# Plant attributes interact with fungal pathogens and nitrogen addition to drive soil enzymatic activities and their temporal variation

**DOI:** 10.1101/2022.09.29.510102

**Authors:** Thu Zar Nwe, Nadia I. Maaroufi, Eric Allan, Santiago Soliveres, Anne Kempel

## Abstract

- Nitrogen enrichment can alter soil communities and their functioning directly, via changes in nutrient availability and stoichiometry, or indirectly, by changing plant communities or higher trophic levels. In addition, soil biota and their associated functions may show strong temporal changes in their response to environmental changes, yet most current studies have only focused on one of these potential drivers or have measured soil functioning only once during the peak growing season. Therefore, we know little about the relative importance of the different mechanisms by which nitrogen enrichment affects soil communities, functioning and temporal stability.
- In a large grassland experiment manipulating nitrogen enrichment, plant species richness, functional composition and foliar pathogen presence, we measured activities of two enzymes, β-glucosidase and acid phosphatase, as indicators of soil functioning. We did so across different seasons and years to assess their temporal dynamics and how consistently they responded to multiple drivers.
- Nitrogen addition was the most important driver of β-glucosidase activity, and it increased β-glucosidase activity over time. However, interactions between plant attributes and fungicide application were the main drivers of acid phosphatase activity. The temporal stability of soil enzyme activity was differently affected by two facets of plant diversity (species richness [+] and functional diversity [-]), with nitrogen and fungicide addition dampening these effects.
- Synthesis: Fungicide effects, and their interactions with plant diversity, show the importance of foliar pathogens not only for above-but also for belowground processes, and highlight the possibility that these plant enemies are major modulators in the relationships between plant diversity and ecosystem functioning. We also show the need to consider temporal dynamics in belowground processes to better understand the responses of ecosystem functioning to environmental changes such as nutrient enrichment.

## Introduction

Nitrogen (N) enrichment from agriculture and atmospheric deposition has increased dramatically since the 1960s and is profoundly altering many aspects of ecosystem functioning. These include the availability, cycling and stoichiometry of different soil nutrients, and the composition and activity of the microbes and fauna living in the soil (Battye et al., 2017; Vitousek et al., 1997; Zhang et al., 2018). Soil extracellular enzymes are a measure of the activity of soil microbiota, and play an important role in key element cycles such as carbon (C) and phosphorus (P) cycles (Sinsabaugh et al., 2008). Soil enzymatic activities depend on resource levels and on microbial diversity and composition (Schimel & Schaeffer, 2012; Sinsabaugh et al., 2008, 2009; Vitousek, 2004), which can be affected directly and indirectly by N addition. Nitrogen can directly affect enzymatic activities by enhancing their production (as it is a major element in their synthesis; (Vitousek, 2004), or by altering soil nutrient levels, and their stoichiometry, and therefore the microbial demand for nutrients. An increase in soil N would be expected to increase C cycling rates, as these two biogeochemical cycles are very closely linked (e.g., Delgado-Baquerizo et al., 2013). N addition might also affect the activity of enzymes linked to P cycling, as it is expected to reduce soil N:P ratios and thereby increase P limitation. Although P availability is mainly determined by soil parent material composition and age, the microbial demand for this nutrient, and therefore P-related soil enzymatic activities, can increase under P limitation (Cline et al., 2018). In addition to these direct effects of N on enzymatic activity and nutrient cycling, a range of indirect effects, mediated by changes in plant and consumer communities, are also possible.

N enrichment could indirectly affect enzymatic activities by altering the diversity and functional composition of the plant community (Cline et al., 2018; Eisenhauer et al., 2010, 2013; Grigulis et al., 2013). The loss of plant diversity following N addition is well known (e.g. Stevens et al., 2004) and several experiments manipulating plant diversity have shown that microbial communities are less active when diversity is reduced (Eisenhauer et al., 2010) and that enzymatic activities vary with diversity (Weisser et al., 2017). Diversity can affect enzymatic activities by altering the quantity of litter entering the soil, as diverse communities are typically more productive than less diverse ones (e.g. Marquard et al., 2009), or by altering the diversity of resources entering the soil (Steinauer et al., 2016). A high litter diversity has been shown to increase decomposition in some cases (Handa et al., 2014; Le Bagousse-Pinguet et al., 2021) and an increase in plant diversity might similarly increase microbial activity and therefore enzyme production. In addition to reducing plant diversity, N addition shifts plant community composition towards dominance by plant species with a fast and acquisitive resource use strategy (Lavorel & Grigulis, 2012). The resource economics spectrum is a major axis of plant functional variation that distinguishes slow growing species, that are conservative in their resource use, from fast growing species that are able to rapidly acquire resources. It is indicated by several key traits, including leaf nutrient contents and specific leaf area (Díaz et al., 2016). This change in plant functional composition has effects on many aspects of biogeochemical cycling, as a shift towards faster plant species is associated with a shift to bacterial dominated systems and faster rates of decomposition and nutrient cycling, which could alter enzymatic activities (de Vries et al., 2012; Grigulis et al., 2013). However, separate effects of diversity and composition are challenging to explore without factorial experiments, as they are often correlated in nature (e.g., Stevens et al. 2004; Marquard et al. 2009).

Nitrogen addition might also increase the abundance of plant enemies such as foliar fungal pathogens (Mitchell et al., 2003) and this could further alter nutrient cycling and enzymatic activities. Few studies have focused on the importance of foliar pathogens in altering enzymatic activities specifically, but pathogens have been shown to alter litter decomposition (Lemons et al., 2005; Wolfe & Ballhorn, 2020). Pathogen exclusion might also alter the effects of diversity on functioning: soil pathogens can drive a positive effect of diversity on biomass production (Maron et al., 2011) and removal of foliar fungal pathogens might therefore dampen effects of diversity on enzymatic activities if monocultures perform better when their specialist pathogens are removed. However, the importance of these various indirect effects (mediated by plant diversity, functional composition or foliar pathogens), relative to the direct effect of N addition, in changing enzymatic activities remains unknown.

Soil microbial communities are very dynamic in time and their abundance and activity vary seasonally (Eisenhauer et al., 2018; Habekost et al., 2008; Regan et al., 2014). These changes could result in seasonal variation in the direct and indirect effects of N on enzymatic activities (but see Siebert et al., 2019 for consistent effects across seasons). For example, pathogen-or plant-mediated mechanisms could be especially relevant during the wettest period or after the peak growing seasons, respectively (e.g., Habekost et al., 2008), in contrast, we might expect a stronger role for direct effects of N addition before the onset of plant growth, just after winter. These seasonal differences could alter the relative importance of different mechanisms in determining soil enzymatic activities throughout the year. The temporal stability of enzymatic activities might therefore vary with N addition, plant diversity and composition, and pathogen abundance, however, this has rarely been explored (Araújo et al., 2013; Habekost et al., 2008; Siebert et al., 2019). In general, intra-annual stability has rarely been explored but consistent functioning across seasons is also likely to be important in generating overall high functioning. A diverse plant community might contain species that have their main period of growth at different times (Allan et al., 2011; Tilman et al., 2006), leading to more consistent C inputs to the soil and therefore more stable enzymatic activity across time. Alternatively, stability can be provided by particular species with highly stable abundance over time (Thibaut & Connolly, 2013) and slow growing species might be expected to provide more consistent inputs to the soil, in comparison to fast growing species that acquire sources rapidly. Periodic inputs of nutrients would be expected to destabilize enzymatic activity by leading to pulses of nutrients. To test these direct and indirect effects of N enrichment on the intra annual variation in enzymatic activities, it is necessary to repeat sampling across different seasons. This repeated sampling would allow us to assess changes in the relative importance of these different factors throughout the year, and not just during the peak growing season.

Here we studied the direct and indirect effects of N on seasonal variation in two soil enzymatic activities related to the C (β-glucosidase) and P (acid phosphatase) cycles. We used an experiment that manipulates plant species richness, functional diversity and composition, together with N addition and foliar pathogen exclusion (Fig. S1). We addressed the following questions: 1) which are the most important drivers of soil enzymatic activities and do these drivers interact? 2) Are these drivers consistent for enzymes related to the C and P cycles? 3) Which factors lead to higher stability of enzyme activities across the year?

## Materials and Methods

### Study site

The study was carried out in the PaNDiV grassland experiment (47°1’18.3”N, 7°27’1.3”E) in Münchenbuchsee, near Bern, Switzerland (Pichon et al., 2020). The grassland was previously extensively managed for sheep grazing and forage production (Cappelli et al., 2020). The soil at the site is a Brunisol (FAO, Cambisol) developed on ground moraine deposits. The background N deposition rate in the area is approximately 17.5 kg N ha^−1^ year^−1^ (Rihm & Künzle, 2019). Mean annual precipitation and temperature at the site are approximately 1021 mm and +9.4°C, respectively.

### Experimental design

In autumn 2015, we set up a field experiment with factorial manipulations of plant species richness, plant functional composition, N enrichment, and foliar fungal pathogens. The experiment consisted of 216 plots of 2m x 2m separated by a 1m buffer zone and arranged in four blocks. The 20 plant species that were used to assemble the plant communities consisted of herbs and grasses that are common in lowland central European grasslands (Cappelli et al., 2020). These species were categorised as those with a fast-growing, acquisitive resource use strategy (referred to as fast species), and those with a slow-growing, conservative strategy (referred to as slow species), based on two plant traits: specific leaf area (SLA), and leaf N content; high values of both indicate faster species (Díaz et al., 2016; Reich, 2014).

To manipulate species richness, we established plots with 1, 4, 8 or all 20 species. To manipulate the composition and diversity of fast/slow functional traits, the plots with four and eight species could contain either only fast, only slow, or a mix of fast and slow species. This allowed us to create a large gradient in functional composition (mean SLA) and functional diversity (in terms of the variance in SLA), independent of species richness. We established all 20 monoculture plots and four replicates of the 20 species plots. For the four and eight species plots we randomly selected five species compositions for each functional composition (fast, slow and mixed) to give 15 combinations of four and 15 of eight species. Each of the 54 different species compositions (20 monocultures + 15 4-spp combinations + 15 8-spp combinations + 4 20-spp replicates) then received the four combinations of N enrichment (no N [control] vs. addition of 100 kg N ha^−1^ y^−1^) and fungicide application (spraying fungicide to remove foliar fungal pathogens vs a control treatment, spraying only water). The 100 kg N ha^−1^ were added in the form of urea twice a year (early April and late June), which corresponds to fertilization rates in intermediately intensively managed temperate grasslands in Europe (Blüthgen et al. 2012). Fungicide application consisted of spraying fungicide (Score Profi, 24.8 % Difenoconazol 250 g.L^-1^ and Ortiva, 32.8% Chlorothalonil 400 g.L^-1^ 6.56% Azoxystrobin 80 g.L^-1^, Syngeta Agro) four times a year: early April and June, late July and September. The control for the fungicide treatment consisted of spraying the same amount of water at the same times.

Overall, our experimental design resulted in a total of 216 plots. The experimental field was divided into four blocks, each block containing all 54 compositions but with the particular N x fungicide treatment for that composition randomly allocated per block. To maintain species compositions, the plots were weeded three times a year in April, July and September, and mown twice annually in June and August, which corresponds to relatively extensive management in lowland grasslands. Further details about the site characteristics and experimental setup of the PaNDiv Experiment can be found in Fig. S1 and in Pichon et al. (2020).

### Soil sampling

To assess temporal variation in microbial activity and available N and P concentrations (NH_4_^+^, NO_3_^-^ and PO_4_^3-^) in the soil, soil samples were taken in autumn (October) 2017, when the maximum amount of litter enters the soil, and four times during the growing season: at the start (April), at peak biomass before the first cut (in May) and twice between the first and second cut of the grassland (July and August) 2018. Within each plot, soil cores were taken from two random locations, avoiding plot edges. At each location, soil was sampled to 20 cm depth using a 1.5 cm stainless steel soil corer. Soil samples were immediately placed in sealed bags and transported to the laboratory in cooling boxes. Soil samples were sieved with a 2 mm sieve to remove stones and roots. From each soil core, a subsample of 5 g was dried at 105°C for 24 hours to estimate the moisture. To assess soil carbon concentrations we took soil samples of 440 cm^3^ at 25 cm depth in autumn 2017 (just one sampling time). We took two samples per plot, homogenised them and removed stones and living material (roots, fauna). We weighed the samples before and after drying at 65°C for 48 hours.

### Nutrient measurements

We measured the available N and P concentrations (NH_4_^+^, NO_3_^-^ and PO_4_^3-^) in the soil using CaCl_2_ with a Continuous Flow Optical-Absorption Spectrometer (CF-OAP; model Skalar Scan+; Skalar Analytical, Breda, The Netherlands). Analyses of soil total C were conducted in a CNS Analyser at the Institute of Geography of the University of Bern. This allowed us to analyse changes in N:P ratios, C:N ratios and C:P ratios in response to N fertilisation, and also to relate these changes to plant and microbial demand for nutrients (as indicated by the soil enzymatic activities, see below).

### Extracellular enzymatic activities

The seasonal variation in the activities of two hydrolytic enzymes, the β-1,4-glucosidase (BG, EC 3.2.1.21) and the acid phosphatase (AP, EC 3.1.3.2), were assessed by colorimetry according to the method of Tabatabai (1994). The β-1,4-glucosidase is involved in the degradation of cellulose and is mainly produced by soil fungi (Stott et al., 2010). The acid phosphatase is involved in the mineralization of organic phosphate compounds and is produced by plants and soil microbes (Cabugao et al., 2017). We considered measuring other important soil enzymatic activities, such as urease or leucine aminopeptidase (degradation of different N sources), but this proved logistically challenging due to the more complex methodologies involved, and the number of samples and sampling periods that we wanted to analyse. In addition, some of these enzymatic activities are strongly correlated with beta-glucosidase (Sinsabaugh et al. 2008; Delgado-Baquerizo et al. 2013), the results of which we present here. Therefore, although incomplete, we believe our selection of enzymatic activities covers the functional responses of both plants and soil biota to our treatments while providing basic information on the cycles of some of the main organic (carbon) and mineral (phosphorous) nutrients.

In brief, the assays of β-glucosidase and acid phosphatase were conducted by homogenising 0.5 g of soil sample, 2 mL TRIS buffer (pH 6.5) and 0.5 mL 25mM substrates *para*-nitrophenyl-β-D-glucopiranoside (*p*NPG) and *para*-nytrophenyl phosphate (*p*NPP), respectively. Soil slurries were incubated for one hour at 37°C. After the incubation period, 0.5 mL of CaCl_2_l and 2mL of NaOH (0.5 M) were added to the soil slurries, vortexed and centrifuged at 3500 rpm during 5 min. Supernatants were transferred to a 96-well plate and the absorbance was measured with a microplate spectrophotometer (BioTek Instruments, Epoch, SN-1510235) at 400 nm. The concentration of *p*-nytrophenol released by the enzymatic activity was obtained from the absorbance values using *p*-nytrophenol standard curves (0-1000 mg L^-1^). The enzymatic activity was expressed in µmol *p*-nytrophenol (*p*NP) per gram dry soil and incubation time (µmol g^-1^ h^-1^). These values were compared with controls, i.e., the same mixtures (soil + reactives), but where the substrate was added after the incubation period, to account for anything that could affect the absorbance levels other than the enzymatic activity *per se*.

### Plant trait measurements

To characterize functional composition, we calculated the community weighted mean (CWM) of specific leaf area (SLA). The experiment was designed using functional groups (fast vs. slow) but we intended to create a continuous gradient in functional composition and diversity and therefore used trait means and variance in the analysis. We measured SLA (SLA= leaf area/dry weight, m^2^ ·kg^-1^) in all monocultures in June 2018, August 2017 and August 2018, by sampling one leaf from five individuals per plant species, and measuring leaf area and dry weight, following the protocol of Garnier et al. (2001). We measured plant species abundance by visually estimating the percentage cover of our target species in each plot. We then calculated a CWM SLA value for each plot by multiplying each species’ relative abundance by the mean SLA value of the species in monoculture in the respective treatment, to account for intraspecific trait variation due to the N and fungicide addition. To obtain a measure of functional diversity we calculated the mean pairwise distance (MPD) in SLA between species within communities in each plot (Pichon et al., 2020). We used unweighted MPD, based on the presence/absence of fast and slow species in each plot, as this is least correlated with CWM SLA and other treatments.

### Statistical analysis

We calculated the mean enzymatic activity over time for β-glucosidase and acid phosphatase, and the standard deviation in enzymatic activity, as a measure of variability per plot across all sampling times. We tested how the mean and standard deviation responded to our experimental treatments using linear mixed effect models (lme4 package in R; Bates et al., 2015). As fixed factors we included N addition (yes, no), fungicide application (yes, no), species diversity (1, 4, 8, or 20 species), CWM SLA (functional composition of fast/slow traits) and MPD in SLA (functional diversity of fast/slow traits). We also included all possible two-way and three-way interactions between terms, except for interactions between CWM SLA and MPD SLA (as these are inevitably correlated given that maximum MPD SLA can only occur at intermediate CWM SLA). We included block (4 levels) and species combination (54 levels, i.e., the specific set of species in the plot) as random terms. We also analysed nutrient ratios, N:P, C:P and C:N, using the same model.

We then analysed each sampling time separately, to determine whether the effects of the experimental treatments on the enzymatic activity of β-glucosidase and acid phosphatase varied over time, using the same models as for the means. In the models for April, May and July we further tested whether the response of enzymatic activities to the experimental treatments and plant community characteristics depended on the N:P ratio in the soil, by adding N:P ratio as a covariate to the models (we only had N:P ratio data for these sampling times). In addition, we tested whether the averaged N:P ratio across the year, the C:N and the C:P ratio affected the enzymatic activities by adding each ratio to our models for the mean enzyme activities (C was measured only once per year). All continuous variables were standardised to a mean of zero and a standard deviation of 1, so we could compare their effects. We derived significances using likelihood-ratio tests, comparing models with and without the factor of interest. Model estimates and 95 % confidence intervals were obtained with the *effects* package (Fox, 2003). All statistical analyses were carried out in R (R Core Team 2018).

## Results

We found contrasting drivers of the two soil enzymatic activities. The mean acid phosphatase was driven by plant attributes and their interaction with fungicide application, while β-glucosidase was not significantly affected by any of the experimental treatments but did tend to be higher with N addition.

Acid phosphatase activity was strongly affected by plant community attributes and their interactions with fungicide. The mean acid phosphatase activity was affected by plant species richness and community weighted mean SLA, however the effect depended on fungicide application (species richness x SLA x fungicide interaction, Table S2, Figure 1a), particularly in October and April (Fig. S3, Table S4). Without fungicide application, acid phosphatase activity slightly increased with species richness and CWM SLA, and the effects of species diversity were consistently positive during four out of five sampling periods. When fungal pathogens were excluded by spraying fungicide, acid phosphatase activity was reduced in fast-growing plant communities, at high plant diversity. In contrast, fungicide application increased acid phosphatase in fast growing monocultures. The mean acid phosphatase activity was also higher in plots with high mean pairwise distance in SLA (MPD SLA; Figure 1b, Table S2). Nitrogen addition did not affect the mean acid phosphatase activity.

**Figure 1.**
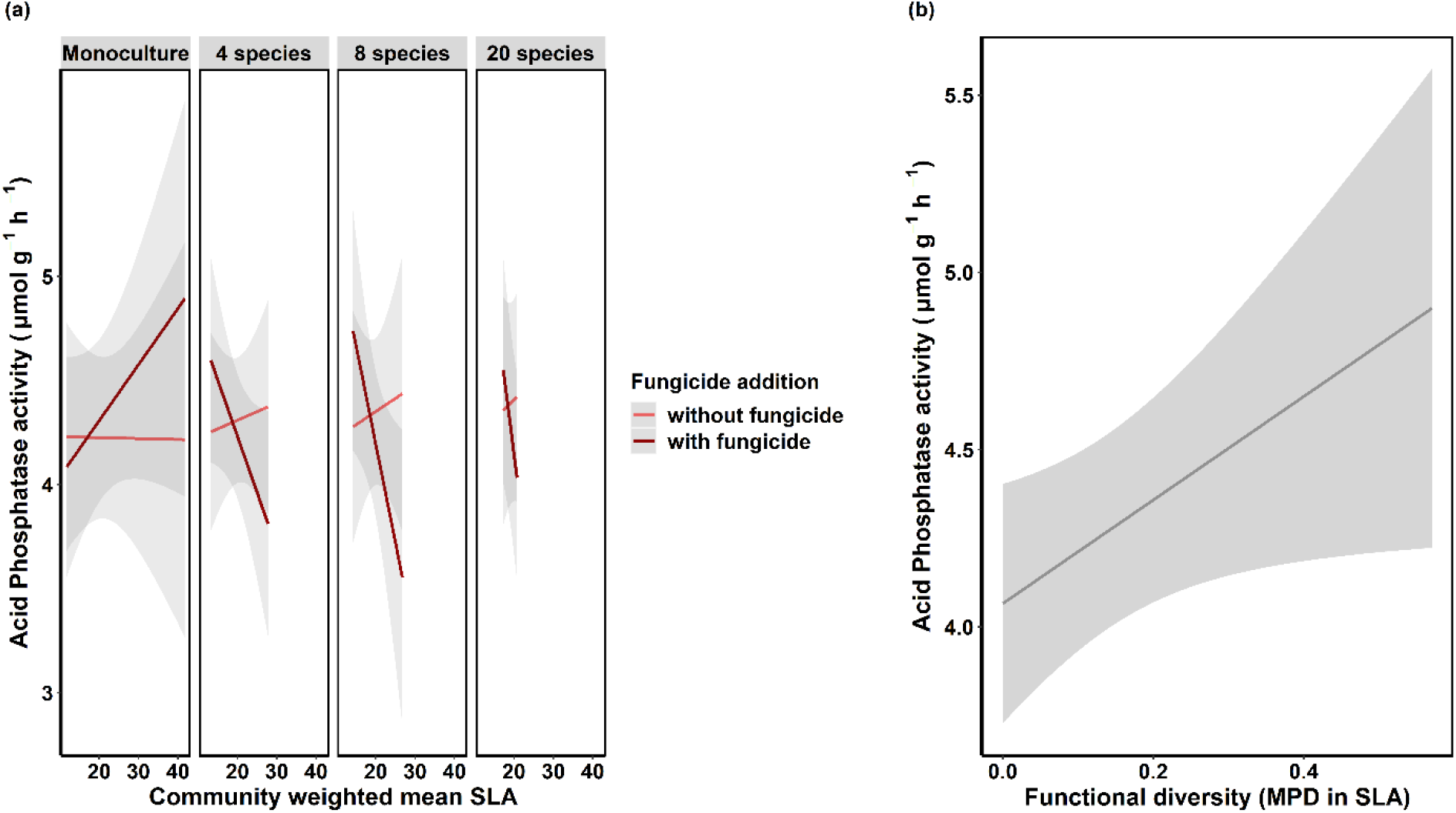
a) Effects of plant species richness, plant community weighted mean SLA and fungicide application and b) plant functional diversity (MPD in SLA) on mean acid phosphatase activity. We show fitted values from a generalised linear mixed effect model (Table S1). Shaded areas represent 95 % confidence intervals. Confidence intervals were derived from the effect package.

The mean β-glucosidase activity over all five sampling times tended to be higher in fertilised plots (marginally significant N effect, Table S3, Figure 2). However, no other factors affected mean β-glucosidase activity (Table S3). Analysing the different sampling periods revealed that in April and May 2018, during the peak growing season, plant attributes interacted with N and fungicide addition to determine β-glucosidase activity and fungicide had effects on β-glucosidase in some seasons, often in interaction with plant attributes (Figure S4, Table S5).

**Figure 2.**
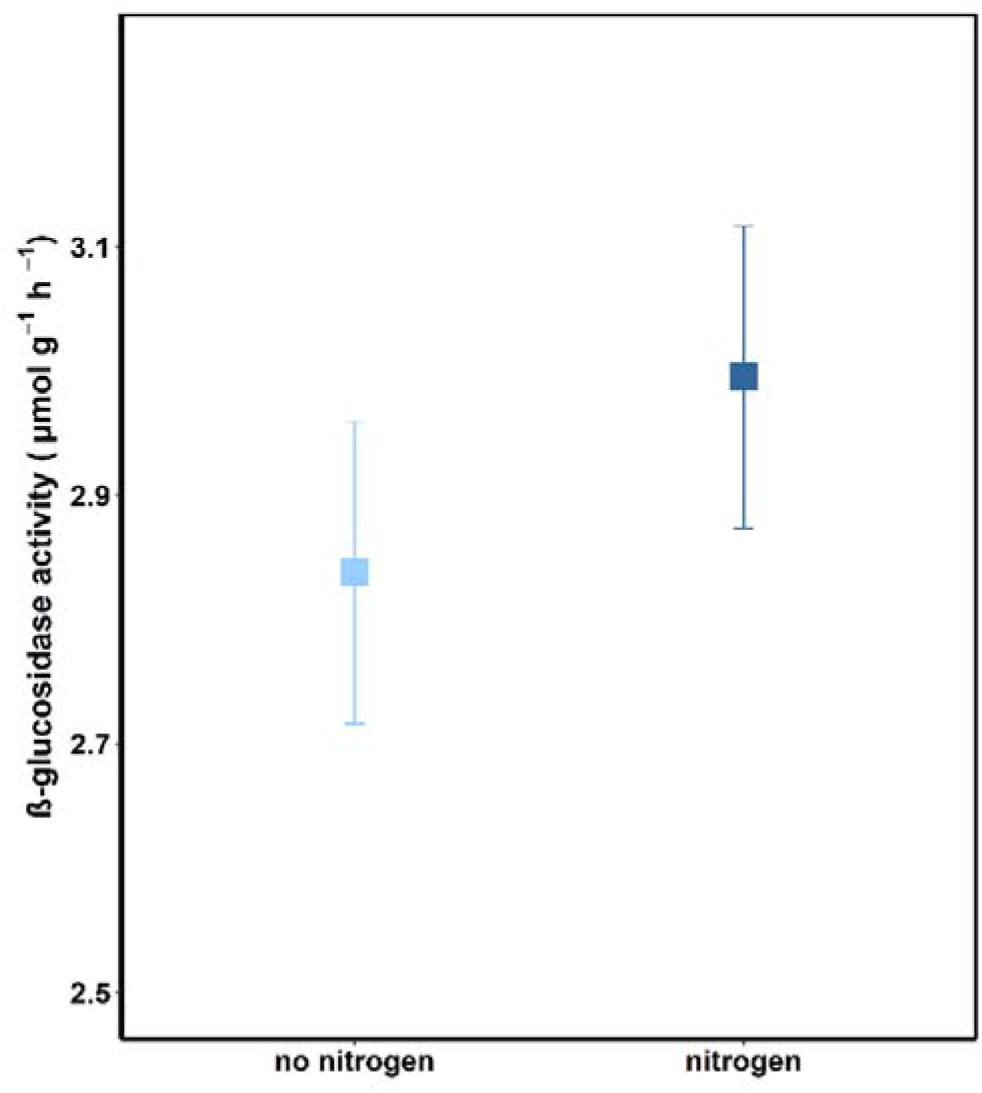
Effects of nitrogen application on mean β-glucosidase activity. We show fitted values and 95 % confidence interval from a generalized linear mixed effect model (Table S2). Confidence intervals were derived from the effects package.

When we tested how the different variables affected the stability of enzymatic activities across time, we found that the stability of enzyme activities was affected differently by the two facets of plant diversity (plant species richness and functional diversity), and that N addition, or the removal of fungal pathogens, dampened the effect of diversity (Figures 3-4, Table S2, S3). The variation in acid phosphatase activity (standard deviation across the year) decreased strongly with plant taxonomic richness, and therefore stability was higher in communities with high richness. However, when we added N or sprayed fungicide, the positive effect of plant species richness on stability disappeared or was reversed (species richness x fungicide x N interaction, Table S2, Figure 3). In contrast, the temporal variation in both enzyme activities was highest in plant communities with a high functional diversity, indicating that functional diversity destabilized enzymatic activities (Table S2, S3, Figure 4). The negative effect of MPD SLA on enzyme stability disappeared when we added N or fungicide alone (MPD SLA x fungicide x N interaction, Table S2, S3).

**Figure 3.**
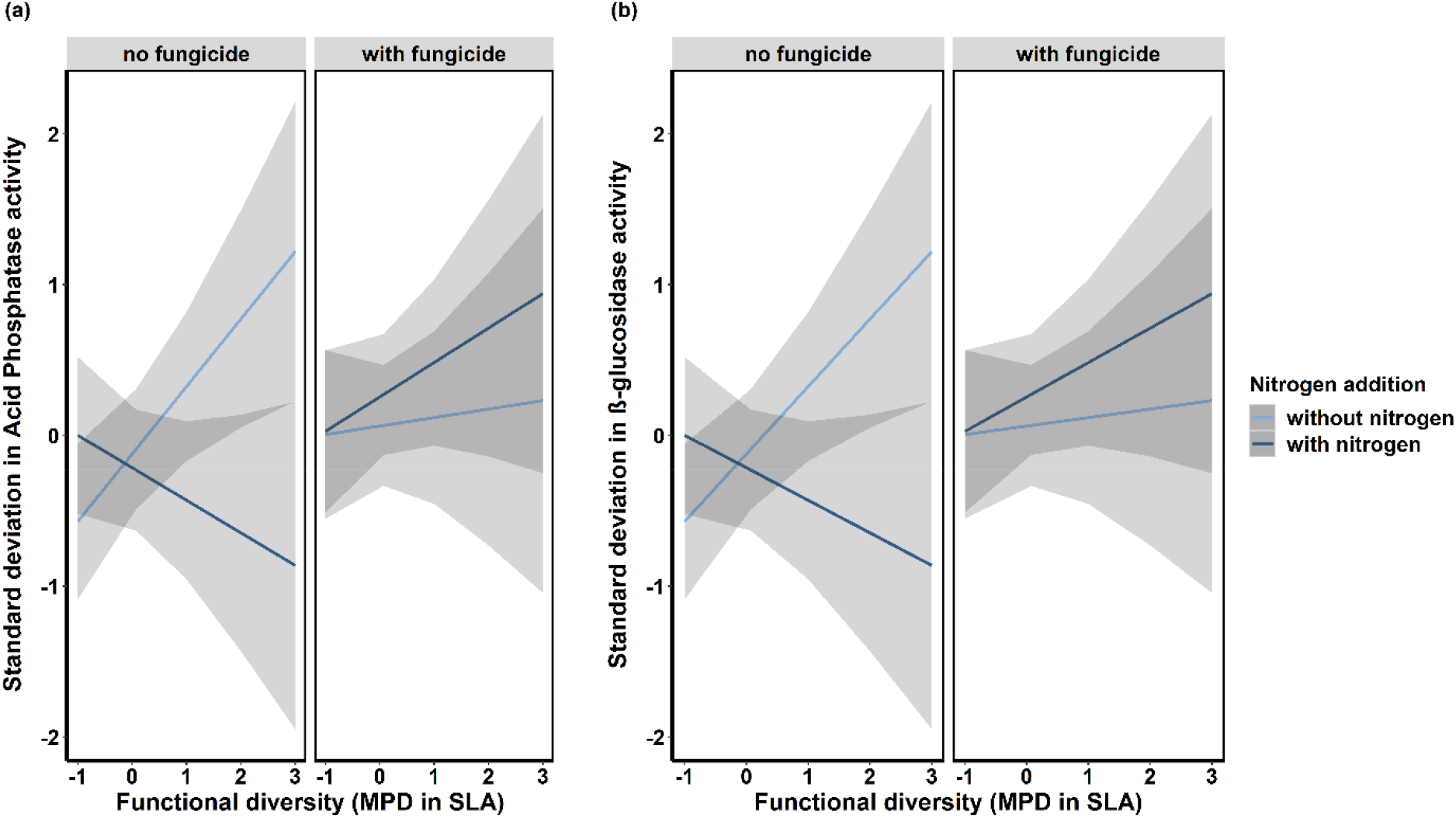
Effects of functional diversity, nitrogen addition and fungicide application on the temporal variation of (a) phosphatase activity and (b) β-glucosidase activity. We show fitted values from a generalized linear mixed effect model (Table S1-S2). Shaded areas represent 95 % confidence intervals. Confidence intervals were derived from the effect package.

**Figure 4.**
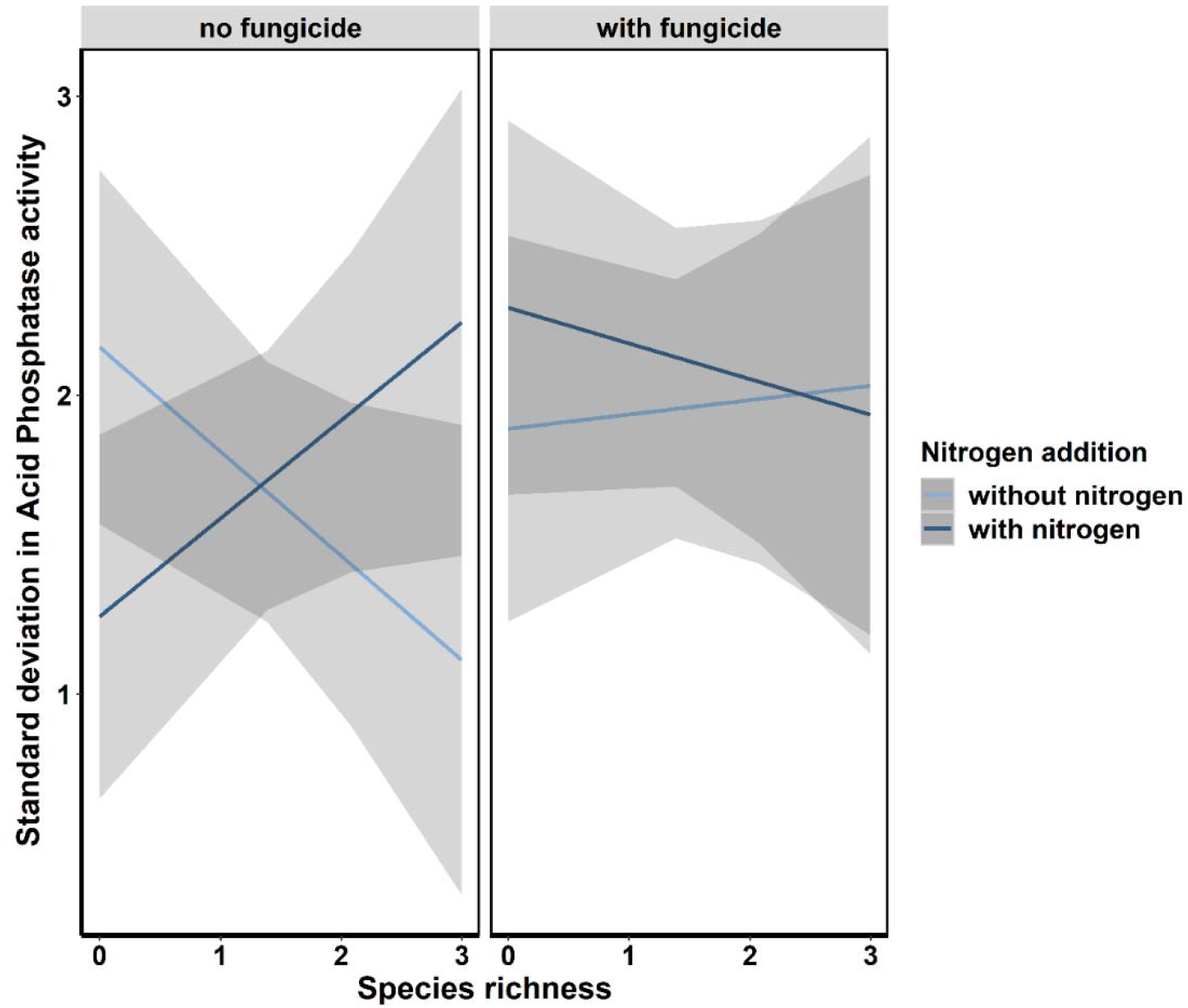
Effects of plant species richness, nitrogen addition and fungicide application on the temporal variation of acid phosphatase activity. We show fitted values from a generalized linear mixed effect model (Table S1). Shaded areas represent 95 % confidence intervals. Confidence intervals were derived from the effect package.

For three of the five time points (April, May, July), we also tested whether the soil N:P ratio affected the response of β-glucosidase and acid phosphatase activity to the experimental treatments and the plant community characteristics. Although the addition of N significantly increased the N:P ratio in April and May and showed the same tendency in July 2018 (Figure S5, Table S6), these changes in N:P ratio did not affect any of the enzymatic activities (Table S8, S9). We also tested whether the yearly average of the N:P ratio, the C:N ratio or the C:P ratio affected the response of β-glucosidase and acid phosphatase activity to the experimental treatments and the plant community characteristics. We found that the addition of N significantly increased the yearly average in N:P ratio (Table S7) and there was an interaction between nitrogen addition, species richness and SLA. For the C:P ratio, we found an interaction between plant species richness and functional diversity. However, these changes in the ratios did not alter the way the enzyme activities were affected by the experimental treatments and the plant community characteristics (Table S7, S10, S11).

## Discussion

We found that plant diversity was a major driver of both mean and variation in enzymatic activities. However, different diversity dimensions had different effects, and removal of fungal pathogens and addition of N modified the effects of diversity on enzymatic activities. This highlights the importance of better understanding the context dependency of diversity effects on functioning and shows that indirect effects of nitrogen addition can be major drivers of soil enzymatic activities.

### Different drivers of acid phosphatase and β-glucosidase activities

We found different drivers of the activity of β-glucosidase and acid phosphatase. Acid phosphatase enzymatic activities were driven by complex interactive effects of plant functional composition (CWM SLA), diversity (MPD SLA and species richness) and plant enemies (foliar pathogens). In contrast, mean levels of β-glucosidase activity were not significantly affected by any of the experimental treatments, although N had a marginally significant effect. Surprisingly, although N:P ratios did increase with our N addition treatment (Table 1, Table S1), this increase did not affect soil enzymatic activities, as the N:P ratio was not selected in the most parsimonious models explaining either β-glucosidase or acid phosphatase enzymatic activities (Table S6, S7). Acid phosphatase may not have been affected by N:P ratios because N availability remained low (N:P ratio <7) which indicates an N-limited environment, even in fertilized plots (Venterink & Güsewell, 2010). This results highlights the strong homeostasis that soil microbes can show in response to changes in nutrient stoichiometry (Zhan et al., 2017). Our results therefore show that β-glucosidase activity is overall quite resistant to the various direct and indirect effects of N that we simulated in our experiment, however, acid phosphatase activity is affected by interactions between various indirect effects.

**Table 1.** N:P ratios measured in control, N-fertilised (Fertilized, fungicide sprayed (Fungicide) and N-fertilized and fungicide sprayed plots at three different time points. Results show the mean and standard error (*N* = 54).

| | N ($\text{NO}_3^- + \text{NH}_4^+$) : P ($\text{PO}_4^{3-}$) ratio | | | | | |
| --- | --- | --- | --- | --- | --- | --- |
|  | April |  | May |  | July |  |
|  | Mean | SE | Mean | SE | Mean | SE |
| Control | 1.39 | 0.35 | 1.68 | 0.24 | 2.61 | 0.29 |
| Fungicide | 0.53 | 0.12 | 1.18 | 0.14 | 2.48 | 0.26 |
| Fertilized | 2.94 | 0.47 | 2.37 | 0.41 | 3.54 | 0.44 |
| Fertilized + Fungicide | 2.91 | 0.56 | 2.01 | 0.37 | 3.05 | 0.47 |

Acid phosphatase enzymatic activities showed complex responses to our experimental treatments, with significant 3-way and 2-way interactions between plant attributes, N and fungicide application. Phosphatase is secreted by both plant roots and soil fungi (Cabugao et al., 2017), as opposed to β -glucosidase, which is only produced by soil microbes (Zang et al., 2018). The latter could explain why acid phosphatase was strongly determined by plant attributes (richness and functional composition), and their interactions with higher trophic levels (fungicide treatment). For example, fast-growing plant communities with high SLA had lower acid phosphatase activity, but this effect was only evident in the absence of foliar pathogens, and in particular at high plant diversity. One explanation for this might be that plants infested with pathogens release more root exudates to the soil to stimulate the growth of beneficial microbes.

In addition, several foliar pathogens are also facultative saprotrophs, which can feed on dead or decaying organic material (Suzuki & Sasaki, 2019), and they might contribute to stimulating enzymatic activities in the soil when senesced leaves fall as litter. In our experiment, fast-growing plant species showed higher levels of pathogen infection (Cappelli et al., 2020), particularly at high plant diversity. When sprayed with fungicide, foliar pathogens were thus particularly reduced in fast, diverse communities, which may have led to the decrease in enzymatic activity in those communities (Figure 1a, Fungicide x CWM SLA x Plant species richness interaction). Another explanation for the effect of fungicide might be that it directly harmed soil microbial communities. Other studies found negative effects of one of the active ingredient of our fungicide (Difenoconazole) on several enzymatic activities when applied at doses slightly higher than ours (Roman et al., 2021). However, non-target effects of the fungicide would not explain why the negative effect of fungicide on acid phosphatase activity is only evident in fast communities and at high diversity. In contrast, at low diversity and high SLA, fungicide actually increased acid phosphatase activity, which would not be expected if the fungicide affects enzymatic activities by directly killing soil fungi. Also note that low diversity plant communities are those with most bare soil (Cappelli et al., 2020) and we might therefore expect the greatest direct effect of fungicide on the soil microbes at low diversity. Therefore, while we cannot rule out some influence of non-target effects, we feel the more likely explanation is that foliar pathogens play an important role in affecting soil functioning.

Phosphatase activity also increased in communities with a high functional diversity (Jing et al., 2016), which has been previously observed for ecosystem functioning in general and for other facets of nutrient cycling (i.e., litter decomposition; Handa et al., 2014) and also for soil enzymatic activities (e.g., Le Bagousse-Pinguet et al., 2021). More diverse plant assemblages often show complementarity in resource use and biomass growth, which could lead to a higher P demand and therefore a larger release of plant-generated phosphatase. In addition, more functionally diverse communities produce more diverse litter, which can then stimulate decomposition and microbial growth, enhancing the release of microbe-generated phosphatase.

The trend for an increase in β-glucosidase activity following N addition could be caused by an enhanced microbial foraging for soil C (and the associated release of β-glucosidase) derived from a stronger demand for C caused by N addition (Xiao et al., 2018). Although previous studies found variable effects of N addition on β-glucosidase activity (Jing et al., 2016; Zheng et al., 2015 with neutral or negative effects of N on β-glucosidase), our results are in accordance with a recent meta-analysis, which showed increased activity of enzymes related to C-acquisition following N addition (Xiao et al., 2018). For β-glucosidase, our results thus partially support the resource allocation theory of enzyme production, which indicates that nutrient addition increases the activity of nutrient-releasing enzymes (Allison & Vitousek, 2005; Sinsabaugh & Moorhead, 1994).

### Plant diversity is a major driver of intra-annual stability in enzymatic activities

The stability of both soil enzymatic activities increased with plant species richness but decreased with functional diversity, and both effects were dampened when adding N or removing foliar pathogens. While aboveground phenological changes have received considerable research attention (e.g., García-Palacios et al., 2018; Peñuelas & Filella, 2001), those occurring belowground are far less studied, despite their importance for crucial ecosystem processes such as soil C storage (Eisenhauer et al., 2018). By assessing the temporal dynamics of belowground processes, our study showed that the main drivers of soil enzymatic activities varied across the year (Tables S2-S4), which could explain contrasting results in previous literature (e.g., those of N addition on β-glucosidase activity, discussed above). For example, we found contrasting effects of species richness and functional diversity on acid phosphatase activity depending on the sampling period (Tables S4), and similar contrasting effects of N addition and fungicide application on β-glucosidase activity (Table S5) depending on when we sampled. It seems as if species richness and functional diversity increased acid phosphatase activity mainly at peak biomass but had no or slightly negative effects in other times of the year. Similarly, N addition increased β-glucosidase activity only after its application in April and July. The temporal changes in the effects of all of our treatments translated into significant interactions between plant attributes, foliar pathogens and N addition in determining the stability of enzymatic activities across time.

Plant diversity commonly stabilises ecosystem functioning because different species vary in abundance across time (Hautier et al., 2014; Hector et al., 2010). Our finding that plant species richness increased the stability of acid phosphatase activity supports those ideas, and shows that plant species richness also stabilises belowground functions across seasons within a year. However, the positive effect of plant species richness on enzyme stability was dampened or even reversed, when our plant communities were fertilised or sprayed with fungicide. This is similar to findings of Hautier et al. (2014) who showed that the positive effect of plant species richness on the stability of aboveground plant biomass production disappeared when grasslands had been fertilized, because N addition synchronised species fluctuations across years (see also Liu et al., 2019). There are several reasons why N addition and fungicide application could alter diversity-stability relationships: first, the addition of N might lead to pulses of nutrients, directly reducing the stability of acid phosphatase activity by increasing variation in the demand for P. Second, plant species richness might protect plant communities against pathogen driven fluctuations in carbon inputs. Removing pathogens using fungicide might remove the stabilizing effect of species richness.

Although N addition and fungicide application dampened effects of both species richness and functional diversity, these two diversity dimensions had contrasting effects on stability. Plant species richness stabilised soil enzymatic activities through time, whereas functional diversity destabilized enzymatic activities across seasons (see control “no fungicide, no fertilization” plots in Figs. 3 and 4). The negative effects of functional diversity on stability may be driven by the difference in performance between fast and slow-growing plant species in our experiment. Fast-growing species grow early in the season but do not recover after the first cut. Thus, communities with a high functional diversity in SLA show a larger shift in functional composition throughout the year. As fast and slow-growing plant species may differ in their carbon supply rates to the soil, carbon supply may be more variable in functionally diverse communities. Similar to species richness, the effect of functional diversity (MPD SLA) on temporal variation of both enzymes disappeared when we sprayed fungicide or added N (MPD SLA x fungicide x nitrogen interaction, Figure 3, Table S, S3). The addition of N and the removal of fungal pathogens are likely to have allowed fast-growing species to recover later in the season, leading to the coexistence of both slow and fast growing species and a more stable supply rate of carbon to the soil. The combined treatment of N and fungicide, however, led to dominance of fast species and lowered the stability of enzyme activity. These results underline the importance of considering the effects of different diversity dimensions on stability as species richness and functional diversity can have contrasting effects.

## Conclusion

By evaluating several drivers of enzymatic activities, we found that acid phosphatase was driven by plant attributes and their interaction with fungicide application, while β-glucosidase was less responsive and only slightly affected by N addition. The importance of these drivers not only varied with the enzymatic activity, but also across time, with more complex interactions often found when evaluating the stability of enzymatic activities. Our results have several implications. Firstly, the effects of fungicide on soil enzymatic activities suggests that foliar pathogens may be important not only for above, but also for belowground processes, and the interactions of fungicide with plant species richness also highlight that these plant enemies can modulate the relationships between plant diversity and ecosystem functioning. Second, our study highlights the importance of evaluating temporal dynamics in belowground processes, and the relative importance of their different drivers, to fully understand the responses of ecosystem functioning to ongoing global change. Given the importance of soil extracellular enzymes for key element cycles we need to better understand these complex responses.

## Acknowledgements

We thank Marlise Zimmermann, Mervi Laitinen, Hugo Vincent and the members of the Allan Lab for support with field and labwork. We also thank Noémie Anna Pichon for help with the analysis of the data. This work is part of the PaNDiv grassland experiment, which was funded by the Swiss National Science Foundation (Project Number: 310030_185260). We thank the Ramony Cajal programme (RyC-2016-20604) founded by Spanish Ministry of Science, the Swedish Research Council for Sustainable Development for funding NIM (FORMAS 2018-00748), the Swiss National Science Foundation for funding AK (P3P3PA_160992), and the Federal Commission for Scholarships for foreign students (FCS) for funding TZN.

