## Supporting information for "Plant attributes interact with fungal pathogens and nitrogen addition to drive soil enzymatic activities and their temporal variation"

**Supplementary results:**

Overall treatments, acid phosphatase activity was highest in April and August, and lowest in July and β-glucosidase activity was highest in July and August, and lowest in May (see Figure S1).

We also analysed the effects of all variables on β-glucosidase and acid phosphatase activity at each individual sampling time. For β-glucosidase activity, this showed that the positive effect of N was mainly present over the peak growing season (April, May, July), whereas the negative effect of fungicide on β-glucosidase activity was mainly found early and at the end of the growing season (April, August, October). In April and May, we also found several interacting effects (Figure S1 (a-b), Table S3-S4), however, we do not discuss them further as they only appeared once.

Plant species richness consistently increased acid phosphatase activity, however in April and October the positive effect disappeared and became negative in plots with nitrogen addition or fungicide application. Similarly, in October we found an interacting effect of SLA, nitrogen addition and fungicide application, and in April an interacting effect of SLA and nitrogen. This shows that, in general, acid phosphatase activity is higher in plots with fast growing plant communities, but that nitrogen addition and fungicide application strongly reduces acid phosphatase activity in fast-growing communities, reversing the pattern (Figure S1 (a-b), Table S3-S4).

**Supplementary Figures**

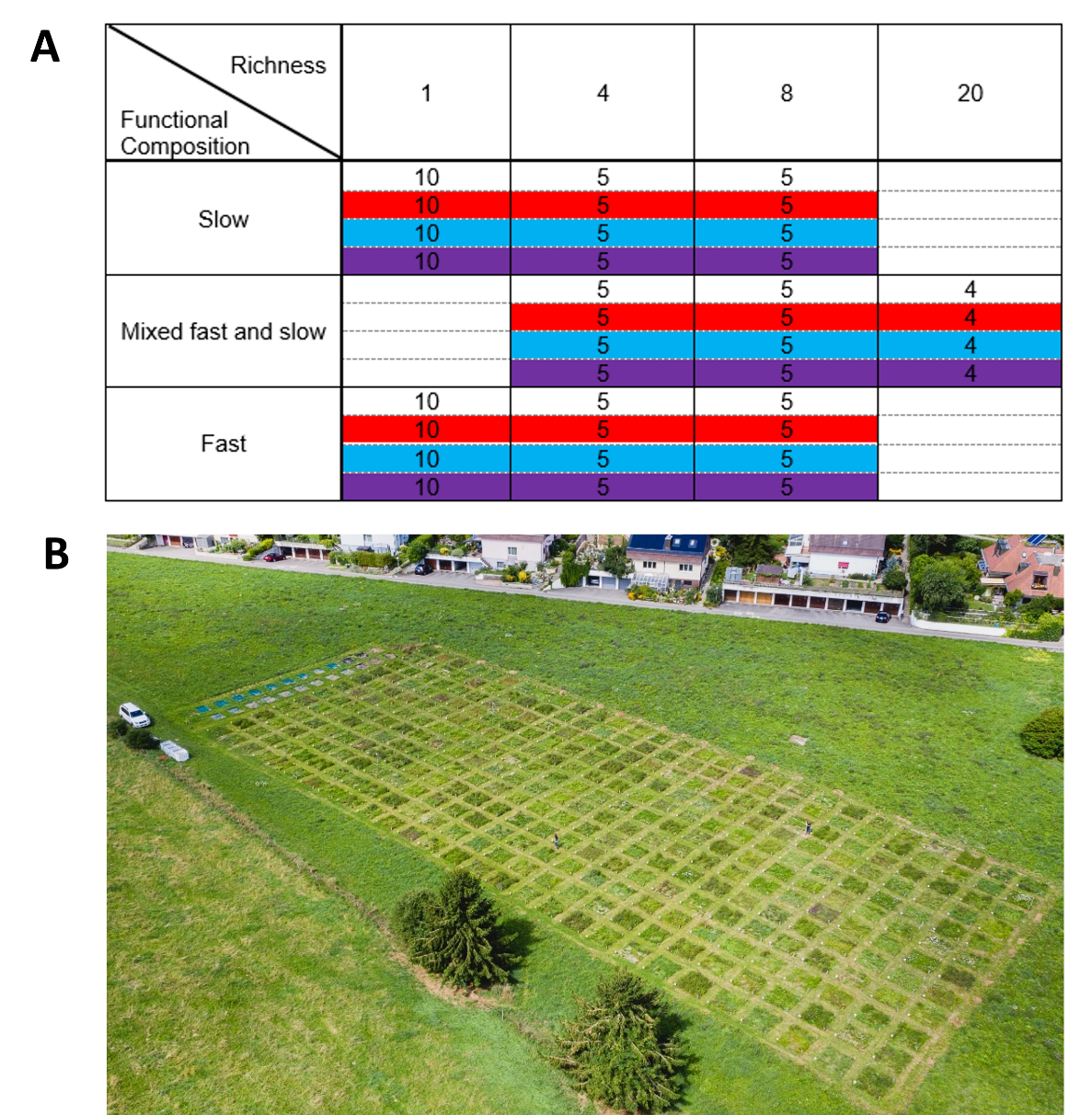

Fig. S1. A) The PaNDiv experimental design. The numbers in the cells show the number of replicates of each combination of species richness, functional composition and the four treatments. The experiment contains four species richness levels, monocultures (1 species) and mixtures of 4, 8 and 20 species. In addition, there are three functional compositions, i.e., plots with only slow, only fast or a mix of fast and slow species. The nitrogen (Urea 100 kg ha^-1^y^-1^) x fungicide (Score Profi [24.8%] with active ingredient Difenoconazol [250 g·L-1] and Ortiva [32.8%] with active ingredients Chlorothalonil [6.56%] and Azoxystrobin [80 g · L-1]) treatments are shown with colours: white no treatment, red only fungicide, blue only nitrogen and purple nitrogen and fungicide. Note that each species composition is present in every block and receives a different treatment in each block, so there are 5 unique combinations of four slow species and each of these five receive the four treatments. At the 20 species level there is only one combination of species replicated four times. B) aerial view of the PaNDiv Experiment.

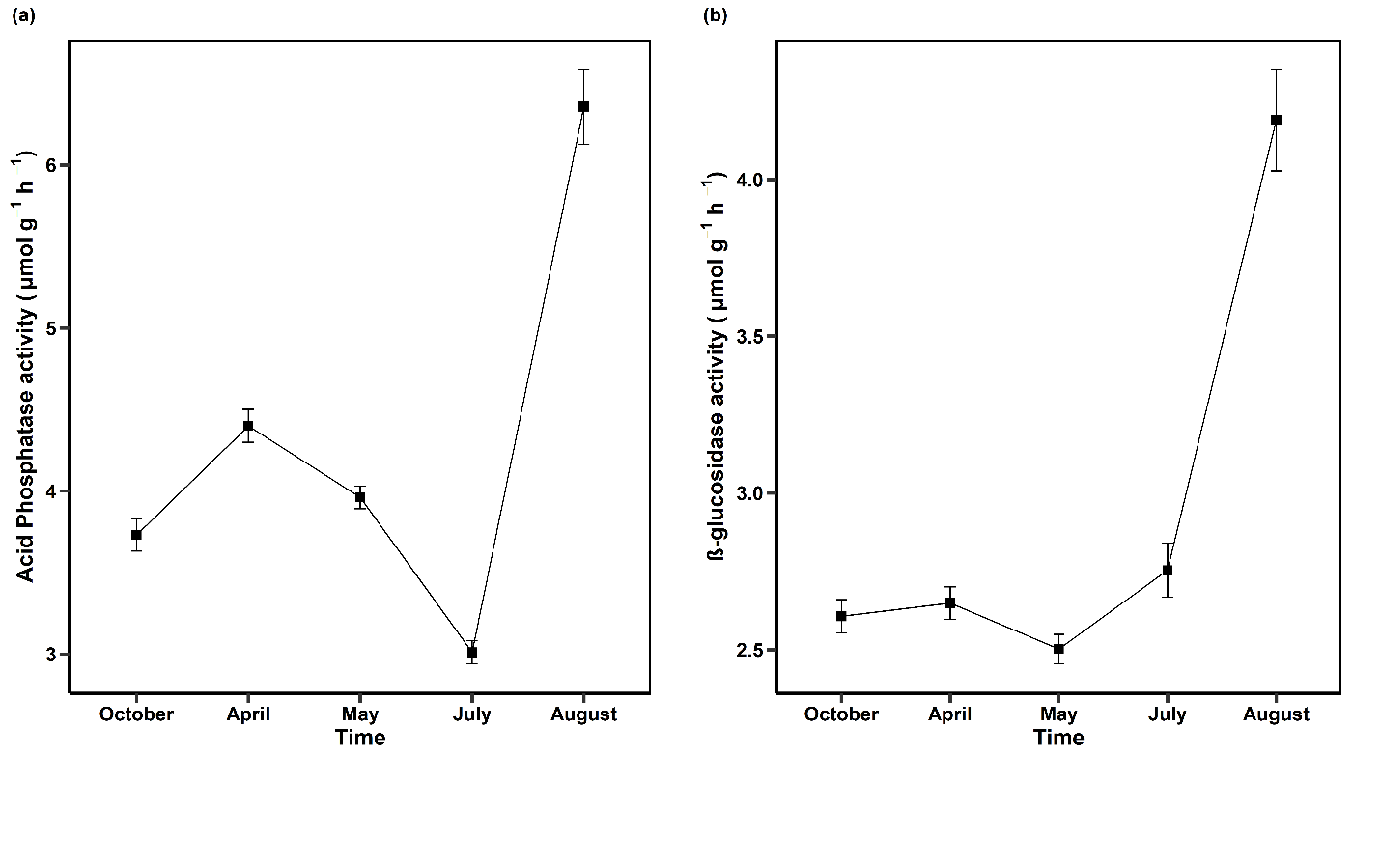

Figure S2. Mean ± standard errors of a) acid phosphatase and b) β-glucosidase activities at the different sampling dates averaged across all treatments.

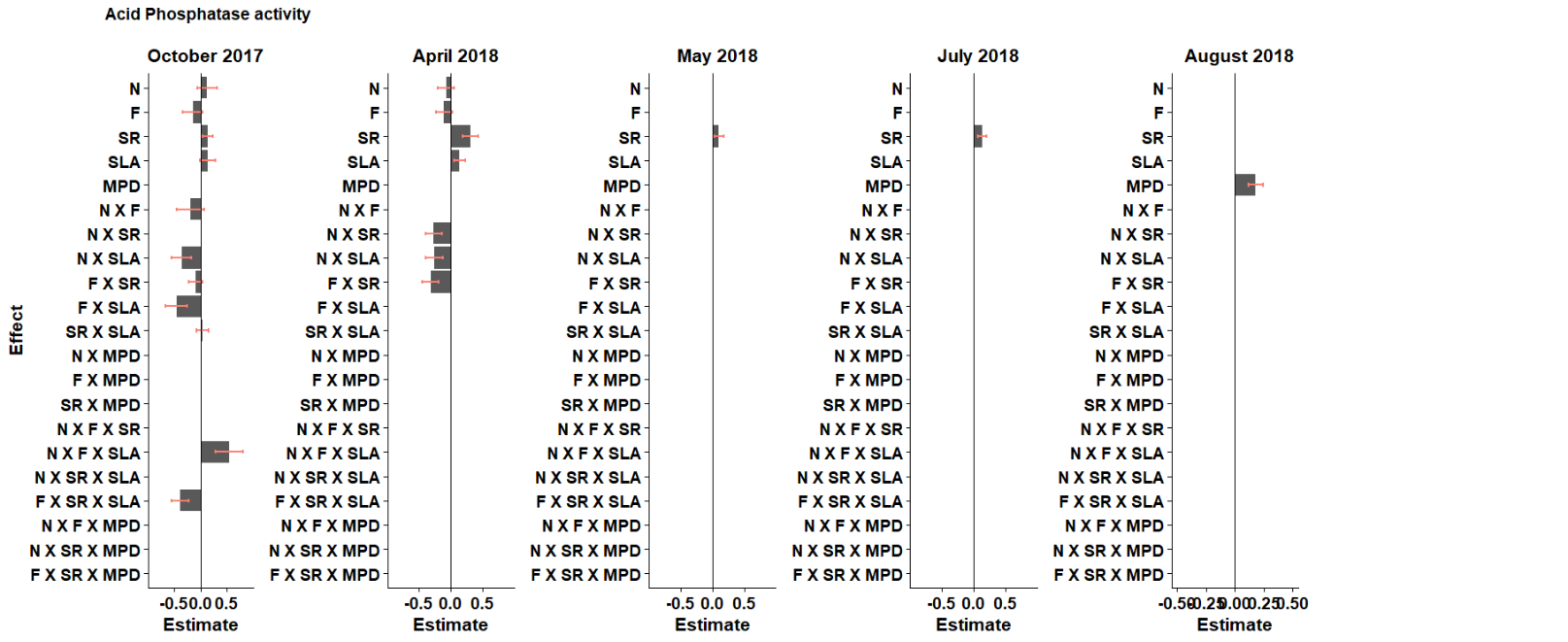

Figure S3. The effect of nitrogen addition (N), fungicide application (F), plant species richness

(SR), plant functional composition (SLA), plant functional diversity (MPD in SLA) and their interactions on acid phosphatase activity at five different time points.

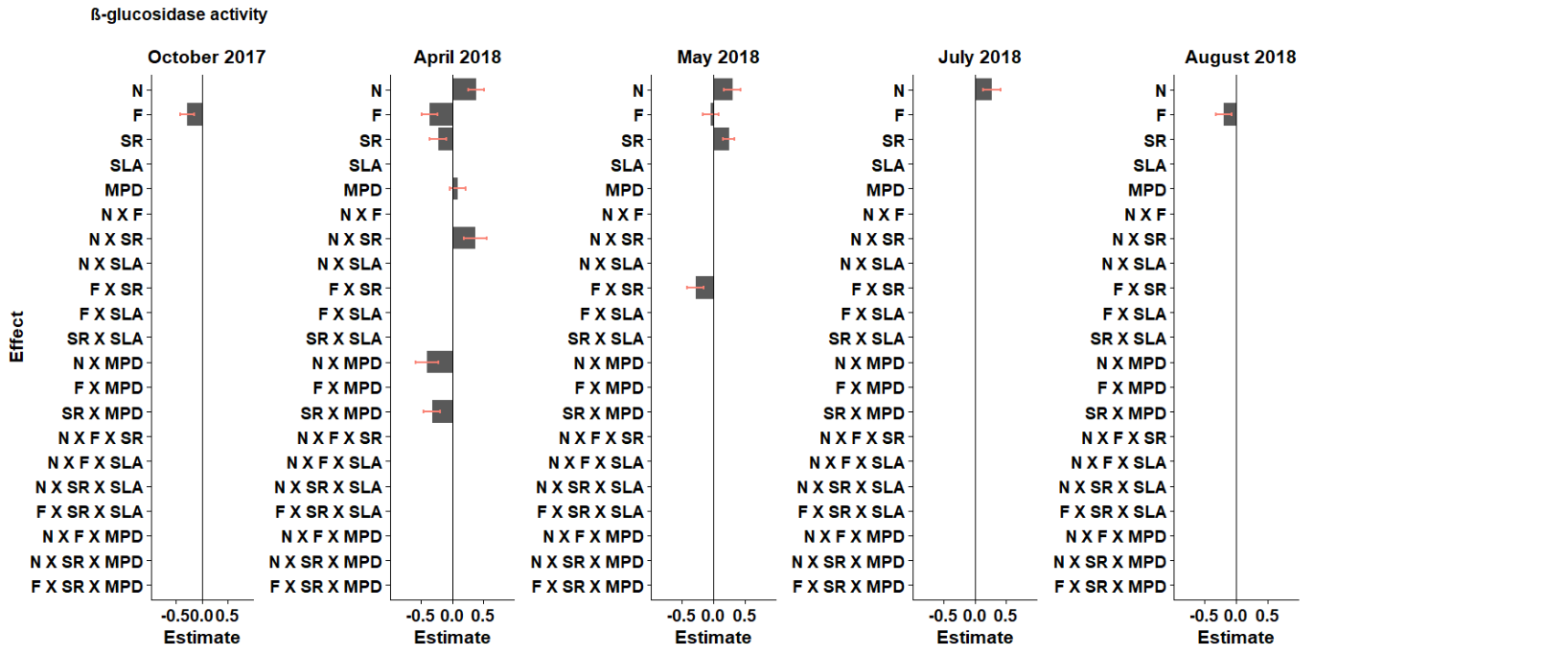

Figure S4. The effect of nitrogen addition (N), fungicide application (F), plant species richness (SR), plant functional composition (SLA), plant functional diversity (MPD in SLA) and their interactions on β-glucosidase activity at five different time points.

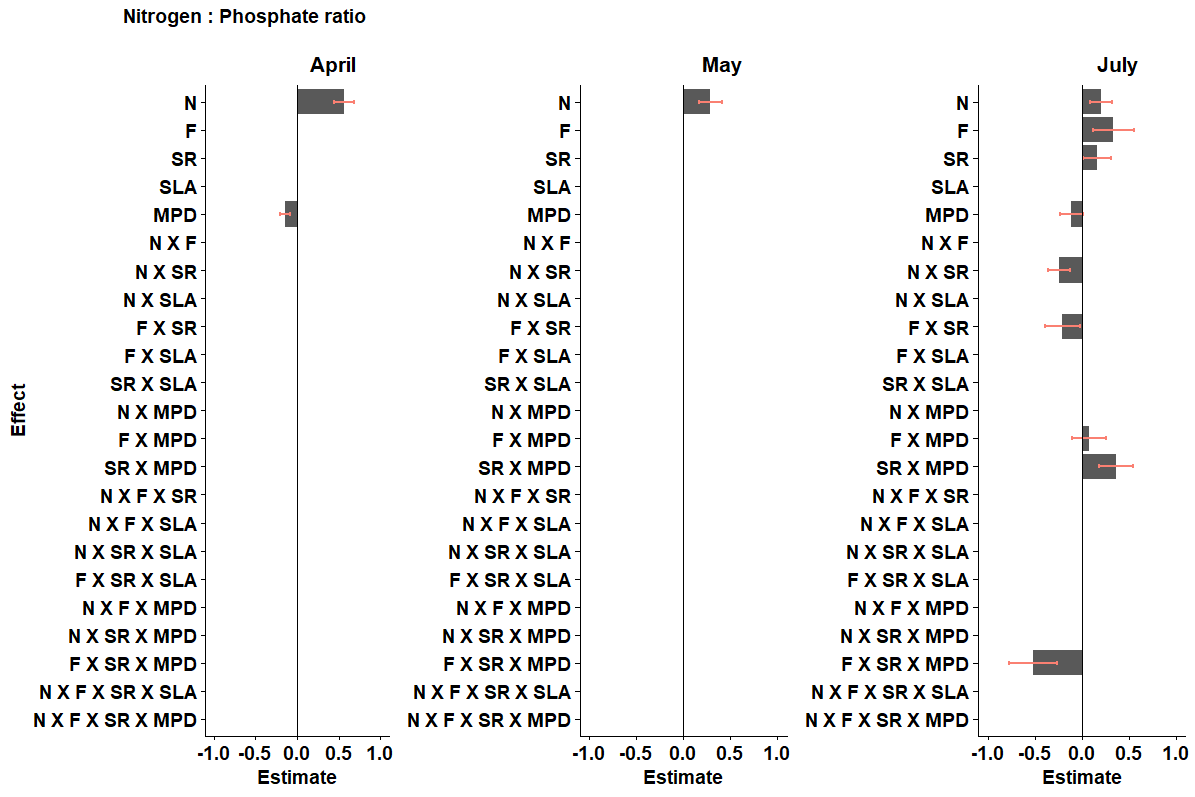

Figure S5. The effect of nitrogen addition (N), fungicide application (F), plant species richness (SR), plant functional composition (SLA), plant functional diversity (MPD in SLA) and their interactions on the N:P ratio measured at three different time points.

**Supplementary tables**

Table S1: C:N ratios, N:P ratios, and C:P ratios measured in control plots, N-fertilised (Fertilized) plots, fungicide treated (Fungicide) plots and plots receiving fertilizer and fungicide. Shown are the means +/-SE (*N* = 54) of the averaged ratios across the year.

| **Treatment** | **C:N ratio** | | **N:P ratio** | | **C:P ratio** | |
| --- | --- | --- | --- | --- | --- | --- |
|  | **Mean** | **SE** | **Mean** | **SE** | **Mean** | **SE** |
| **Control** | 0.29 | 0.04 | 1.69 | 0.25 | 0.32 | 0.03 |
| **Fungicide** | 0.38 | 0.07 | 1.13 | 0.14 | 0.26 | 0.02 |
| **Fertilized** | 0.21 | 0.02 | 2.62 | 0.36 | 0.34 | 0.03 |
| **Fertilized + Fungicide** | 0.32 | 0.08 | 2.41 | 0.4 | 0.34 | 0.03 |

Table S2. Effects of nitrogen addition (N), fungicide application (F), plant species richness (SR), plant functional composition (SLA), plant functional diversity (MPD in SLA) and their interactions on the mean and the stability (measured as standard deviation) of acid phosphatase activity. We used linear mixed models and stepwise removed non-significant terms from the models. We obtained significance by using likelihood-ratio tests comparing models with and without the factor of interest.

| **Acid Phosphatase activity** | | | | | | | | | |
| --- | --- | --- | --- | --- | --- | --- | --- | --- | --- |
| **Factor** | **Coefficient of variation** | | | **Mean** | | | **Standard deviation** | | |
|  | **Estimate** | **SE** | **P-value** | **Estimate** | **SE** | **P-value** | **Estimate** | **SE** | **P-value** |
| **(Intercept)** | -0.21 | 0.16 | marginal | 0.03 | 0.16 | marginal | -0.12 | 0.20 | marginal |
| **N** | -0.15 | 0.27 | 0.87 | 0.01 | 0.27 | 0.17 | -0.09 | 0.17 | marginal |
| **F** | **0.43** | **0.13** | **< 0.001 **** | -0.08 | 0.13 | marginal | 0.19 | 0.17 | marginal |
| **SR** | -0.36 | 0.19 | 0.37 | 0.07 | 0.12 | marginal | -0.33 | 0.17 | marginal |
| **SLA** | 0.06 | 0.16 | 0.87 | 0.04 | 0.12 | marginal | 0.10 | 0.16 | 0.82 |
| **MPD** | 0.28 | 0.17 | 0.47 | **0.20** | **0.10** | **0.045 *** | 0.45 | 0.16 | marginal |
| **N X F** | 0.29 | 0.25 | 0.33 | 0.23 | 0.25 | 0.38 | 0.29 | 0.24 | marginal |
| **N X SR** | 0.65 | 0.26 | 0.31 | 0.39 | 0.26 | 0.36 | 0.63 | 0.24 | marginal |
| **N X SLA** | 0.15 | 0.23 | 0.72 | -0.21 | 0.23 | 0.41 | 0.03 | 0.22 | 0.81 |
| **F X SR** | 0.28 | 0.28 | 0.93 | -0.16 | 0.13 | marginal | 0.37 | 0.26 | marginal |
| **F X SLA** | 0.02 | 0.22 | 0.67 | -0.25 | 0.17 | marginal | -0.15 | 0.21 | 0.85 |
| **SR X SLA** | 0.15 | 0.14 | 0.38 | 0.03 | 0.11 | marginal | 0.18 | 0.14 | 0.91 |
| **N X MPD** | -0.56 | 0.24 | 0.15 | -0.53 | 0.24 | 0.09 | -0.66 | 0.23 | marginal |
| **F X MPD** | -0.26 | 0.28 | 0.58 | -0.26 | 0.28 | 0.59 | -0.39 | 0.26 | marginal |
| **SR X MPD** | -0.14 | 0.23 | 0.36 | 0.08 | 0.23 | 0.70 | -0.16 | 0.22 | 0.24 |
| **N X F X SR** | -0.85 | 0.38 | 0.33 | -0.65 | 0.37 | 0.10 | **-0.79** | **0.36** | **0.028 *** |
| **N X F X SLA** | -0.42 | 0.26 | 0.11 | -0.03 | 0.26 | 0.91 | -0.26 | 0.25 | 0.32 |
| **N X SR X SLA** | -0.01 | 0.18 | 0.94 | -0.15 | 0.18 | 0.43 | -0.08 | 0.17 | 0.64 |
| **F X SR X SLA** | -0.15 | 0.18 | 0.37 | **-0.33** | **0.16** | **0.043 *** | -0.31 | 0.17 | 0.08 |
| **N X F X MPD** | 0.84 | 0.38 | 0.06 | 0.70 | 0.38 | 0.45 | **0.84** | **0.36** | **0.022 *** |
| **N X SR X MPD** | 0.02 | 0.27 | 0.95 | 0.08 | 0.27 | 0.78 | 0.02 | 0.26 | 0.94 |
| **F X SR X MPD** | -0.02 | 0.27 | 0.94 | -0.13 | 0.27 | 0.64 | 0.00 | 0.27 | 0.99 |

Table S3. Effects of nitrogen addition (N), fungicide application (F), plant species richness (SR), plant functional composition (SLA), plant functional diversity (MPD in SLA) and their interactions on the mean and the stability (measured as standard deviation) of β-glucosidase activity. We used linear mixed models and stepwise removed non-significant terms from the models. We obtained significance by using likelihood-ratio tests comparing models with and without the factor of interest.

| **β-glucosidase activity** | | | | | | | | | |
| --- | --- | --- | --- | --- | --- | --- | --- | --- | --- |
| **Factor** | **Coefficient of variation** | | | **Mean** | | | **Standard deviation** | | |
|  | **Estimate** | **SE** | **P-value** | **Estimate** | **SE** | **P-value** | **Estimate** | **SE** | **P-value** |
| **(Intercept)** | 0.12 | 0.22 | marginal | -0.12 | 0.18 | marginal | 0.12 | 0.24 | marginal |
| **N** | -0.27 | 0.17 | marginal | 0.24 | 0.13 | 0.060^(^*^)^ | -0.18 | 0.17 | marginal |
| **F** | -0.08 | 0.17 | marginal | -0.21 | 0.13 | 0.095^(^*^)^ | -0.19 | 0.16 | marginal |
| **SR** | 0.05 | 0.19 | 0.70 | -0.15 | 0.19 | 0.97 | -0.17 | 0.18 | 0.90 |
| **SLA** | -0.09 | 0.16 | 0.29 | -0.17 | 0.16 | 0.79 | -0.08 | 0.15 | 0.49 |
| **MPD** | 0.31 | 0.11 | marginal | 0.45 | 0.17 | 0.18 | 0.45 | 0.11 | marginal |
| **N X F** | 0.22 | 0.24 | marginal | -0.02 | 0.25 | 0.92 | 0.25 | 0.23 | marginal |
| **N X SR** | -0.19 | 0.25 | 0.32 | 0.52 | 0.26 | 0.09^(^*^)^ | 0.13 | 0.25 | 0.83 |
| **N X SLA** | 0.06 | 0.22 | 0.25 | 0.25 | 0.22 | 0.14 | 0.14 | 0.21 | 0.91 |
| **F X SR** | 0.01 | 0.27 | 0.56 | 0.10 | 0.28 | 0.12 | 0.21 | 0.27 | 0.45 |
| **F X SLA** | -0.03 | 0.21 | 0.47 | 0.41 | 0.22 | 0.38 | 0.04 | 0.21 | 0.55 |
| **SR X SLA** | -0.12 | 0.14 | 0.77 | 0.00 | 0.14 | 0.37 | -0.05 | 0.13 | 0.70 |
| **N X MPD** | -0.47 | 0.16 | marginal | -0.70 | 0.24 | 0.06^(^*^)^ | -0.57 | 0.16 | marginal |
| **F X MPD** | -0.49 | 0.17 | marginal | -0.49 | 0.28 | 0.87 | -0.58 | 0.16 | marginal |
| **SR X MPD** | 0.03 | 0.22 | 0.28 | 0.11 | 0.22 | 0.82 | 0.01 | 0.21 | 0.21 |
| **N X F X SR** | 0.10 | 0.36 | 0.79 | -0.47 | 0.37 | 0.27 | -0.19 | 0.35 | 0.56 |
| **N X F X SLA** | 0.19 | 0.25 | 0.45 | -0.23 | 0.25 | 0.38 | 0.11 | 0.24 | 0.65 |
| **N X SR X SLA** | 0.37 | 0.17 | 0.05 | -0.06 | 0.18 | 0.77 | 0.25 | 0.17 | 0.19 |
| **F X SR X SLA** | -0.05 | 0.17 | 0.78 | 0.24 | 0.17 | 0.18 | 0.02 | 0.17 | 0.89 |
| **N X F X MPD** | **0.80** | **0.24** | **< 0.001 **** | 0.59 | 0.37 | 0.38 | **0.81** | **0.23** | **< 0.001 **** |
| **N X SR X MPD** | -0.22 | 0.26 | 0.37 | 0.07 | 0.27 | 0.79 | -0.22 | 0.25 | 0.35 |
| **F X SR X MPD** | -0.17 | 0.26 | 0.44 | -0.44 | 0.27 | 0.24 | -0.17 | 0.26 | 0.55 |

Table S4: The effect of nitrogen addition (N), fungicide application (F), plant species richness (SR), plant functional composition (SLA), plant functional diversity (MPD in SLA) and their interactions on acid phosphatase activity at the five different time points. We used linear mixed models and stepwise removed non-significant terms from the models. We obtained significance by using

likelihood-ratio tests comparing models with and without the factor of interest.

| **Acid Phosphatase activity** | | | | | | | | | | | | | | | |
| --- | --- | --- | --- | --- | --- | --- | --- | --- | --- | --- | --- | --- | --- | --- | --- |
| **Factor** | **October** | | | **April** | | | **May** | | | **July** | | | **August** | | |
|  | **Estimate** | **SE** | **P-value** | **Estimate** | **SE** | **P-value** | **Estimate** | **SE** | **P-value** | **Estimate** | **SE** | **P-value** | **Estimate** | **SE** | **P-value** |
| **(Intercept)** | 0.04 | 0.13 | marginal | 0.11 | 0.12 | marginal | 0.00 | 0.12 | marginal | 0.00 | 0.08 | marginal | 0.01 | 0.19 | marginal |
| **N** | 0.13 | 0.18 | marginal | -0.07 | 0.13 | marginal | 0.05 | 0.27 | 0.23 | 0.32 | 0.29 | 0.21 | -0.09 | 0.27 | 0.63 |
| **F** | -0.14 | 0.19 | marginal | -0.11 | 0.13 | marginal | -0.21 | 0.27 | 0.47 | 0.13 | 0.29 | 0.89 | -0.15 | 0.28 | 0.67 |
| **SR** | 0.13 | 0.09 | marginal | 0.30 | 0.12 | marginal | 0.11 | 0.08 | 0.19 | **0.14** | **0.07** | **0.042 *** | -0.18 | 0.22 | 0.51 |
| **SLA** | 0.14 | 0.15 | marginal | 0.13 | 0.09 | marginal | 0.13 | 0.17 | 0.77 | 0.02 | 0.17 | 0.88 | 0.13 | 0.17 | 0.67 |
| **MPD** | 0.21 | 0.18 | 0.67 | -0.01 | 0.18 | 0.52 | -0.02 | 0.18 | 0.87 | 0.17 | 0.20 | 0.98 | **0.18** | **0.07** | **0.018 *** |
| **N X F** | -0.19 | 0.26 | marginal | 0.15 | 0.25 | 0.62 | 0.10 | 0.25 | 0.73 | -0.04 | 0.27 | 0.86 | 0.36 | 0.26 | 0.21 |
| **N X SR** | 0.26 | 0.28 | 0.84 | **-0.27** | **0.13** | **0.035 *** | 0.15 | 0.26 | 0.42 | 0.00 | 0.29 | 0.64 | 0.40 | 0.29 | 0.24 |
| **N X SLA** | -0.36 | 0.19 | marginal | **-0.26** | **0.13** | **0.045 *** | 0.01 | 0.23 | 0.64 | 0.06 | 0.25 | 0.65 | -0.07 | 0.23 | 0.85 |
| **F X SR** | -0.10 | 0.13 | marginal | **-0.31** | **0.13** | **0.015 *** | -0.27 | 0.29 | 0.46 | 0.03 | 0.31 | 0.44 | 0.07 | 0.31 | 0.97 |
| **F X SLA** | -0.46 | 0.21 | marginal | 0.10 | 0.22 | 0.58 | -0.25 | 0.22 | 0.48 | -0.18 | 0.24 | 0.12 | -0.22 | 0.23 | 0.85 |
| **SR X SLA** | 0.04 | 0.11 | marginal | 0.05 | 0.15 | 0.24 | 0.07 | 0.15 | 0.69 | -0.05 | 0.15 | 0.37 | 0.17 | 0.15 | 0.65 |
| **N X MPD** | -0.19 | 0.26 | 0.48 | 0.24 | 0.25 | 0.19 | -0.21 | 0.25 | 0.52 | -0.09 | 0.27 | 0.73 | -0.55 | 0.26 | 0.13 |
| **F X MPD** | -0.24 | 0.30 | 0.41 | -0.02 | 0.29 | 0.87 | 0.22 | 0.29 | 0.40 | -0.18 | 0.31 | 0.43 | -0.05 | 0.30 | 0.23 |
| **SR X MPD** | 0.10 | 0.24 | 0.42 | 0.47 | 0.24 | 0.35 | 0.05 | 0.25 | 0.43 | 0.12 | 0.24 | 0.80 | 0.01 | 0.25 | 0.98 |
| **N X F X SR** | -0.29 | 0.39 | 0.44 | -0.05 | 0.37 | 0.89 | 0.07 | 0.38 | 0.79 | 0.14 | 0.41 | 0.75 | -0.57 | 0.39 | 0.17 |
| **N X F X SLA** | **0.54** | **0.26** | **0.041 *** | -0.28 | 0.26 | 0.31 | 0.14 | 0.26 | 0.61 | -0.05 | 0.28 | 0.83 | -0.14 | 0.27 | 0.59 |
| **N X SR X SLA** | 0.12 | 0.19 | 0.50 | -0.29 | 0.18 | 0.23 | 0.03 | 0.18 | 0.87 | -0.04 | 0.20 | 0.86 | -0.17 | 0.18 | 0.37 |
| **F X SR X SLA** | **-0.39** | **0.16** | **0.020 *** | -0.11 | 0.18 | 0.51 | -0.09 | 0.18 | 0.65 | -0.03 | 0.19 | 0.89 | -0.33 | 0.18 | 0.17 |
| **N X F X MPD** | 0.08 | 0.39 | 0.84 | 0.09 | 0.38 | 0.84 | 0.00 | 0.38 | 0.99 | -0.07 | 0.41 | 0.87 | 0.64 | 0.40 | 0.50 |
| **N X SR X MPD** | 0.20 | 0.28 | 0.41 | -0.07 | 0.27 | 0.78 | 0.08 | 0.27 | 0.74 | -0.15 | 0.29 | 0.63 | -0.04 | 0.28 | 0.88 |
| **F X SR X MPD** | -0.20 | 0.28 | 0.48 | -0.49 | 0.27 | 0.06 | 0.08 | 0.27 | 0.77 | -0.19 | 0.29 | 0.49 | 0.02 | 0.28 | 0.95 |

Table S5: The effect of nitrogen addition (N), fungicide application (F), plant species richness (SR), plant functional composition (SLA), plant functional diversity (MPD in SLA) and their interactions on β-glucosidase activity at the five different time points. We used linear mixed models and stepwise removed non-significant terms from the models. We obtained significance by using likelihood-ratio tests comparing models with and without the factor of interest.

| **β-glucosidase activity** | | | | | | | | | | | | | | | |
| --- | --- | --- | --- | --- | --- | --- | --- | --- | --- | --- | --- | --- | --- | --- | --- |
| **Factor** | **October** | | | **April** | | | **May** | | | **July** | | | **August** |  |  |
|  | **Estimate** | **SE** | **P-value** | **Estimate** | **SE** | **P-value** | **Estimate** | **SE** | **P-value** | **Estimate** | **SE** | **P-value** | **Estimate** | **SE** | **P-value** |
| **(Intercept)** | 0.14 | 0.10 | marginal | 0.22 | 0.15 | marginal | -0.13 | 0.13 | marginal | -0.14 | 0.10 | marginal | 0.08 | 0.24 | marginal |
| **N** | -0.36 | 0.28 | 0.83 | 0.38 | 0.13 | marginal | **0.30** | **0.13** | **0.024*** | 0.27 | 0.14 | 0.054^(^*^)^ | -0.01 | 0.26 | 0.76 |
| **F** | **-0.28** | **0.13** | **0.037*** | **-0.36** | **0.13** | **0.006**** | -0.04 | 0.13 | marginal | 0.39 | 0.30 | 0.26 | -0.20 | 0.13 | 0.15 |
| **SR** | 0.22 | 0.20 | 0.80 | -0.22 | 0.14 | marginal | 0.25 | 0.09 | marginal | 0.05 | 0.21 | 0.93 | -0.28 | 0.19 | 0.43 |
| **SLA** | 0.11 | 0.17 | 0.48 | -0.22 | 0.16 | 0.42 | -0.19 | 0.17 | 0.73 | -0.13 | 0.17 | 0.61 | -0.07 | 0.16 | 0.97 |
| **MPD** | -0.08 | 0.18 | 0.81 | 0.09 | 0.13 | marginal | 0.16 | 0.18 | 0.69 | 0.12 | 0.20 | 0.42 | 0.49 | 0.16 | 0.26 |
| **N X F** | -0.15 | 0.26 | 0.65 | -0.25 | 0.25 | 0.37 | -0.19 | 0.26 | 0.48 | -0.05 | 0.27 | 0.89 | 0.23 | 0.25 | 0.30 |
| **N X SR** | 0.05 | 0.28 | 0.93 | **0.37** | **0.19** | **0.048*** | 0.61 | 0.27 | 0.57 | -0.05 | 0.29 | 0.22 | 0.42 | 0.26 | 0.06 |
| **N X SLA** | -0.26 | 0.24 | 0.34 | 0.25 | 0.22 | 0.07 | 0.36 | 0.23 | 0.37 | 0.19 | 0.25 | 0.33 | 0.16 | 0.22 | 0.56 |
| **F X SR** | -0.14 | 0.30 | 0.07 | -0.16 | 0.28 | 0.17 | **-0.29** | **0.13** | **0.031*** | -0.13 | 0.32 | 0.38 | 0.28 | 0.29 | 0.70 |
| **F X SLA** | 0.24 | 0.23 | 0.95 | 0.25 | 0.22 | 0.73 | 0.37 | 0.23 | 0.73 | 0.23 | 0.25 | 0.51 | 0.17 | 0.22 | 0.33 |
| **SR X SLA** | 0.11 | 0.15 | 0.90 | -0.04 | 0.14 | 1.00 | -0.11 | 0.15 | 0.60 | -0.04 | 0.15 | 0.85 | 0.04 | 0.14 | 0.30 |
| **N X MPD** | 0.23 | 0.26 | 0.76 | -0.40 | 0.18 | **0.032*** | -0.35 | 0.25 | 0.21 | -0.28 | 0.27 | 0.26 | -0.67 | 0.23 | 0.15 |
| **F X MPD** | -0.17 | 0.30 | 0.59 | 0.23 | 0.28 | 0.91 | -0.23 | 0.29 | 0.80 | 0.07 | 0.32 | 0.25 | -0.60 | 0.31 | 0.34 |
| **SR X MPD** | 0.13 | 0.24 | 0.05 | **-0.32** | **0.14** | **0.027*** | 0.01 | 0.23 | 0.49 | 0.07 | 0.24 | 0.80 | 0.11 | 0.23 | 0.33 |
| **N X F X SR** | -0.09 | 0.39 | 0.72 | -0.23 | 0.37 | 0.53 | -0.68 | 0.39 | 0.17 | 0.19 | 0.41 | 0.21 | -0.41 | 0.39 | 0.35 |
| **N X F X SLA** | -0.28 | 0.27 | 0.27 | -0.27 | 0.27 | 0.37 | -0.44 | 0.27 | 0.13 | -0.05 | 0.28 | 0.84 | 0.04 | 0.26 | 0.88 |
| **N X SR X SLA** | -0.34 | 0.19 | 0.07 | -0.17 | 0.18 | 0.37 | -0.01 | 0.18 | 0.97 | 0.02 | 0.20 | 0.93 | 0.12 | 0.17 | 0.50 |
| **F X SR X SLA** | 0.11 | 0.18 | 0.58 | 0.25 | 0.18 | 0.15 | 0.30 | 0.18 | 0.12 | 0.12 | 0.20 | 0.57 | 0.03 | 0.17 | 0.86 |
| **N X F X MPD** | 0.00 | 0.39 | 1.00 | -0.31 | 0.38 | 0.08 | 0.38 | 0.39 | 0.31 | 0.15 | 0.41 | 0.73 | 0.66 | 0.40 | 0.14 |
| **N X SR X MPD** | 0.60 | 0.28 | 0.12 | -0.26 | 0.27 | 0.24 | 0.07 | 0.28 | 0.80 | 0.18 | 0.30 | 0.56 | -0.21 | 0.26 | 0.52 |
| **F X SR X MPD** | -0.32 | 0.28 | 0.31 | -0.45 | 0.27 | 0.14 | 0.11 | 0.28 | 0.68 | -0.26 | 0.30 | 0.46 | -0.27 | 0.26 | 0.32 |

Table S6: The effect of nitrogen addition (N), fungicide application (F), plant species richness (SR), plant functional composition (SLA), plant functional diversity (MPD in SLA) and their interactions on the N:P ratio measured at three different time points. We used linear mixed models and stepwise removed non-significant terms from the models. We obtained significance by using likelihood-ratio tests comparing models with and without the factor of interest.

| **N : P ratio** | | | | | | | | | |
| --- | --- | --- | --- | --- | --- | --- | --- | --- | --- |
| **Factor** | **April** | | | **May** | | | **July** | | |
|  | **Estimate** | **SE** | **P-value** | **Estimate** | **SE** | **P-value** | **Estimate** | **SE** | **P-value** |
| **(Intercept)** | -0.28 | 0.18 | marginal | -0.15 | 0.21 | marginal | 2.94 | 0.72 | marginal |
| **N** | **0.56** | **0.12** | **< 0.0001***** | **0.29** | **0.12** | **0.022*** | 1.35 | 0.50 | marginal |
| **F** | 0.00 | 0.32 | 0.38 | -0.07 | 0.32 | 0.37 | -0.38 | 0.52 | marginal |
| **SR** | 0.21 | 0.18 | 0.72 | 0.15 | 0.19 | 0.91 | -0.59 | 0.52 | marginal |
| **SLA** | 0.14 | 0.16 | 0.93 | 0.12 | 0.16 | 0.27 | -0.02 | 0.08 | 0.15 |
| **MPD** | **-0.15** | **0.06** | **0.017*** | -0.17 | 0.17 | 0.06 | -11.47 | 5.38 | marginal |
| **N X F** | -0.19 | 0.44 | 0.18 | 0.39 | 0.44 | 0.42 | -2.93 | 3.47 | 0.78 |
| **N X SR** | -0.33 | 0.25 | 0.53 | 0.08 | 0.26 | 0.53 | **-0.66** | **0.32** | **0.041*** |
| **F X SR** | -0.33 | 0.28 | 0.40 | -0.16 | 0.28 | 0.96 | 0.91 | 0.72 | marginal |
| **N X SLA** | -0.09 | 0.24 | 0.40 | -0.06 | 0.24 | 0.38 | 0.13 | 0.11 | 0.11 |
| **N X MPD** | 0.04 | 0.23 | 0.12 | -0.18 | 0.24 | 0.67 | -1.71 | 10.01 | 0.98 |
| **F X SLA** | -0.17 | 0.23 | 0.80 | -0.18 | 0.24 | 0.64 | -0.01 | 0.11 | 0.72 |
| **F X MPD** | 0.25 | 0.28 | 0.63 | 0.13 | 0.28 | 0.17 | 14.70 | 7.37 | marginal |
| **SR X SLA** | 0.02 | 0.15 | 0.18 | 0.16 | 0.15 | 0.67 | 0.08 | 0.09 | 0.90 |
| **SR X MPD** | 0.29 | 0.24 | 0.88 | 0.31 | 0.25 | 0.24 | 7.73 | 3.83 | marginal |
| **N X F X SR** | 0.42 | 0.39 | 0.87 | -0.26 | 0.40 | 0.86 | -2.48 | 3.83 | 0.80 |
| **N X F X SLA** | 0.47 | 0.37 | 0.34 | 0.21 | 0.38 | 0.78 | 0.10 | 0.16 | 0.29 |
| **N X F X MPD** | -0.54 | 0.39 | 0.23 | 0.26 | 0.40 | 0.47 | 3.54 | 14.36 | 0.13 |
| **N X SR X SLA** | 0.15 | 0.22 | 0.12 | -0.31 | 0.22 | 0.47 | -0.21 | 0.13 | 0.08 |
| **N X SR X MPD** | -0.64 | 0.35 | 0.34 | 0.05 | 0.36 | 0.83 | -1.19 | 7.45 | 0.96 |
| **F X SR X SLA** | -0.05 | 0.21 | 0.69 | -0.23 | 0.22 | 0.90 | -0.04 | 0.13 | 0.96 |
| **F X SR X MPD** | -0.37 | 0.35 | 0.89 | -0.24 | 0.36 | 0.19 | **-11.00** | **5.41** | **0.043*** |
| **N X F X SR X SLA** | 0.28 | 0.34 | 0.40 | 0.43 | 0.35 | 0.11 | 0.10 | 0.20 | 0.56 |
| **N X F X SR X MPD** | 0.68 | 0.51 | 0.10 | -0.22 | 0.52 | 0.67 | 3.05 | 10.84 | 0.78 |

Table S7: The effect of nitrogen addition (N), fungicide application (F), plant species richness (SR), plant functional composition (SLA), plant functional diversity (MPD in SLA) and their interactions on the N:P ratio, C:N ratio and C:P ratio averaged across the year. We used linear mixed models and stepwise removed non-significant terms from the models. We obtained significance by using likelihood-ratio tests comparing models with and without the factor of interest.

| **N : P ratio** | | | | | | | | | |
| --- | --- | --- | --- | --- | --- | --- | --- | --- | --- |
| **Factor** | **N:P ratio** | | | **C:N ratio** | | | **C:P ratio** | | |
|  | **Estimate** | **SE** | **P-value** | **Estimate** | **SE** | **P-value** | **Estimate** | **SE** | **P-value** |
| **(Intercept)** | -0.20 | 0.23 | marginal | 0.07 | 0.16 | marginal | -0.20 | 0.26 | marginal |
| **N** | **0.40** | **0.12** | **0.001**** | -0.12 | 0.13 | marginal | 0.22 | 0.30 | 0.38 |
| **F** | 0.09 | 0.31 | 0.39 | -0.23 | 0.35 | 0.13 | 0.17 | 0.31 | 0.65 |
| **SR** | 0.23 | 0.18 | 0.89 | 0.02 | 0.10 | marginal | 0.26 | 0.09 | marginal |
| **SLA** | 0.16 | 0.15 | 0.21 | 0.02 | 0.12 | marginal | 0.35 | 0.16 | 0.08 |
| **MPD** | **-0.15** | **0.06** | **0.014*** | 0.04 | 0.18 | 0.93 | -0.16 | 0.09 | marginal |
| **N X F** | 0.13 | 0.42 | 0.26 | 0.69 | 0.47 | 0.98 | -0.04 | 0.43 | 0.10 |
| **N X SR** | -0.15 | 0.25 | 0.11 | 0.10 | 0.13 | marginal | -0.38 | 0.25 | 0.75 |
| **F X SR** | -0.27 | 0.27 | 0.66 | 0.15 | 0.30 | 0.29 | -0.53 | 0.27 | 0.61 |
| **N X SLA** | -0.15 | 0.23 | 0.80 | -0.30 | 0.18 | marginal | -0.38 | 0.23 | 0.98 |
| **N X MPD** | -0.17 | 0.23 | 0.45 | -0.03 | 0.25 | 0.74 | -0.06 | 0.23 | 0.15 |
| **F X SLA** | -0.17 | 0.22 | 0.69 | -0.30 | 0.25 | 0.32 | -0.40 | 0.23 | 0.73 |
| **F X MPD** | 0.10 | 0.27 | 0.41 | -0.16 | 0.30 | 0.75 | 0.24 | 0.27 | 0.60 |
| **SR X SLA** | 0.10 | 0.14 | 0.48 | 0.13 | 0.11 | marginal | 0.25 | 0.15 | 0.98 |
| **SR X MPD** | 0.38 | 0.24 | 0.51 | -0.30 | 0.29 | 0.74 | **0.29** | **0.13** | **0.022*** |
| **N X F X SR** | 0.04 | 0.37 | 0.91 | -0.15 | 0.41 | 0.69 | 0.47 | 0.38 | 0.83 |
| **N X F X SLA** | 0.42 | 0.36 | 0.13 | -0.20 | 0.39 | 0.61 | 0.48 | 0.36 | 0.47 |
| **N X F X MPD** | 0.06 | 0.38 | 0.91 | 0.31 | 0.42 | 0.21 | -0.01 | 0.38 | 0.29 |
| **N X SR X SLA** | -0.21 | 0.21 | 0.57 | -0.33 | 0.17 | 0.05^(^*^)^ | -0.37 | 0.22 | 0.24 |
| **N X SR X MPD** | -0.24 | 0.34 | 0.54 | 0.42 | 0.40 | 0.81 | -0.48 | 0.36 | 0.37 |
| **F X SR X SLA** | -0.12 | 0.21 | 0.75 | -0.01 | 0.23 | 0.32 | -0.35 | 0.21 | 0.22 |
| **F X SR X MPD** | -0.46 | 0.34 | 0.13 | 0.58 | 0.40 | 0.89 | -0.67 | 0.36 | 0.09 |
| **N X F X SR X SLA** | 0.42 | 0.33 | 0.13 | -0.40 | 0.36 | 0.27 | 0.38 | 0.33 | 0.26 |
| **N X F X SR X MPD** | 0.19 | 0.49 | 0.69 | -0.93 | 0.55 | 0.07 | 0.68 | 0.50 | 0.06 |

Table S8: The main effect of nitrogen addition (N), fungicide application (F), plant species richness (SR), plant functional composition (SLA), plant functional diversity (MPD in SLA) and N:P ratio on the acid phosphatase activity at the three different time points. We used linear mixed models and stepwise removed non-significant terms from the models. We obtained significance by using likelihood-ratio tests comparing models with and without the factor of interest.

| **Acid Phosphatase activity** | | | | | | | | | |
| --- | --- | --- | --- | --- | --- | --- | --- | --- | --- |
| **Factor** | **April** | | | **May** | | | **July** | | |
|  | **Estimate** | **SE** | **P-value** | **Estimate** | **SE** | **P-value** | **Estimate** | **SE** | **P-value** |
| **(Intercept)** | -0.14 | 0.18 | marginal | -0.14 | 0.18 | marginal | -0.01 | 0.15 | marginal |
| **N:P ratio** | -0.13 | 0.07 | 0.058^(^*^)^ | -0.01 | 0.07 | 0.90 | 0.02 | 0.07 | 0.80 |
| **N** | **0.28** | **0.13** | **0.009**** | **0.28** | **0.13** | **0.032*** | 0.16 | 0.13 | 0.21 |
| **F** | -0.17 | 0.13 | 0.18 | -0.16 | 0.13 | 0.20 | -0.04 | 0.13 | 0.74 |
| **SR** | 0.03 | 0.09 | 0.46 | 0.03 | 0.10 | 0.72 | 0.05 | 0.11 | 0.65 |
| **SLA** | 0.03 | 0.06 | 0.58 | 0.03 | 0.07 | 0.69 | 0.01 | 0.07 | 0.83 |
| **MPD** | 0.03 | 0.10 | 0.76 | 0.04 | 0.10 | 0.26 | **0.16** | **0.07** | **0.020*** |

Table S9: The main effect of nitrogen addition (N), fungicide application (F), plant species richness (SR), plant functional composition (SLA), plant functional diversity (MPD in SLA) and N:P ratio on the β-glucosidase activity at three different time points. We used linear mixed models and stepwise removed non-significant terms from the models. We obtained significance by using likelihood-ratio tests comparing models with and without the factor of interest.

| **β-glucosidase activity** | | | | | | | | | |
| --- | --- | --- | --- | --- | --- | --- | --- | --- | --- |
| **Factor** | **April** | | | **May** | | | **July** | | |
|  | **Estimate** | **SE** | **P-value** | **Estimate** | **SE** | **P-value** | **Estimate** | **SE** | **P-value** |
| **(Intercept)** | -0.12 | 0.18 | marginal | -0.14 | 0.18 | marginal | -0.14 | 0.18 | marginal |
| **N:P ratio** | -0.11 | 0.07 | 0.09 | -0.01 | 0.07 | 0.90 | -0.04 | 0.07 | 0.63 |
| **N** | 0.24 | 0.13 | 0.057^(^*^)^ | 0.28 | 0.13 | **0.032*** | 0.28 | 0.13 | **0.032*** |
| **F** | -0.22 | 0.13 | 0.090^(^*^)^ | -0.16 | 0.13 | 0.20 | -0.16 | 0.13 | 0.20 |
| **SR** | 0.00 | 0.09 | 0.96 | 0.03 | 0.10 | 0.72 | 0.03 | 0.09 | 0.76 |
| **SLA** | 0.02 | 0.07 | 0.79 | 0.03 | 0.07 | 0.69 | 0.03 | 0.06 | 0.64 |
| **MPD** | 0.07 | 0.09 | 0.29 | 0.04 | 0.10 | 0.26 | 0.04 | 0.10 | 0.26 |

Table S10: The main effect of nitrogen addition (N), fungicide application (F), plant species richness (SR), plant functional composition (SLA), plant functional diversity (MPD in SLA) and N:P ratio, CN: ratio and C:P ratio – averaged across the year - on the Acid phosphatase activity. We used linear mixed models and stepwise removed non-significant terms from the models. We obtained significance by using likelihood-ratio tests comparing models with and without the factor of interest. Including the ratios in the analysis does not change the effects of the treatments on enzyme activity.

| **Acid Phosphatase activity** | | | | | | | | | |
| --- | --- | --- | --- | --- | --- | --- | --- | --- | --- |
| **Factor** | **N:P ratio** | | | **C:N ratio** | | | **C:P**  **ratio** | | |
|  | **Estimate** | **SE** | **P-value** | **Estimate** | **SE** | **P-value** | **Estimate** | **SE** | **P-value** |
| **(Intercept)** | -0.042 | 0.177 | marginal | -0.023 | 0.178 | marginal | -0.039 | 0.175 | marginal |
| **Ratios** | -0.014 | 0.074 | 0.865 | 0.003 | 0.069 | 0.97 | -0.034 | 0.074 | 0.675 |
| **N** | 0.147 | 0.132 | 0.27 | 0.127 | 0.132 | 0.325 | 0.142 | 0.131 | 0.28 |
| **F** | -0.063 | 0.129 | 0.637 | -0.069 | 0.132 | 0.597 | -0.061 | 0.131 | 0.651 |
| **SR** | 0.042 | 0.095 | 0.667 | 0.054 | 0.097 | 0.6 | 0.055 | 0.098 | 0.654 |
| **SLA** | 0.009 | 0.066 | 0.893 | 0.011 | 0.067 | 0.872 | 0.014 | 0.067 | 0.836 |
| **MPD** | 0.147 | 0.095 | 0.006** | 0.132 | 0.096 | 0.011* | 0.132 | 0.097 | 0.011* |

Table S11: The main effect of nitrogen addition (N), fungicide application (F), plant species richness (SR), plant functional composition (SLA), plant functional diversity (MPD in SLA) and N:P ratio, CN: ratio and C:P ratio – averaged across the year - on the β-glucosidase activity. We used linear mixed models and stepwise removed non-significant terms from the models. We obtained significance by using likelihood-ratio tests comparing models with and without the factor of interest. Including the ratios in the analysis does not change the effects of the treatments on enzyme activity.

| **β-glucosidase a** **ctivity** | | | | | | | | | |
| --- | --- | --- | --- | --- | --- | --- | --- | --- | --- |
| **Factor** | **N:P ratio** | | | **C:N ratio** | | | **C:P ratio** | | |
|  | **Estimate** | **SE** | **P-value** | **Estimate** | **SE** | **P-value** | **Estimate** | **SE** | **P-value** |
| **(Intercept)** | -0.027 | 0.187 | marginal | -0.022 | 0.206 | marginal | -0.026 | 0.187 | marginal |
| **Ratios** | -0.066 | 0.073 | 0.395 | -0.087 | 0.068 | 0.197 | -0.09 | 0.074 | 0.69 |
| **N** | 0.269 | 0.131 | 0.06 | 0.233 | 0.131 | 0.068 | 0.255 | 0.13 | 0.06 |
| **F** | -0.215 | 0.128 | 0.095 | -0.189 | 0.13 | 0.114 | -0.205 | 0.129 | 0.12 |
| **SR** | -0.003 | 0.094 | 0.971 | 0.005 | 0.096 | 0.958 | 0.016 | 0.097 | 0.868 |
| **SLA** | 0.022 | 0.065 | 0.731 | 0.009 | 0.066 | 0.892 | 0.028 | 0.066 | 0.69 |
| **MPD** | 0.076 | 0.094 | 0.178 | 0.079 | 0.095 | 0.214 | 0.065 | 0.096 | 0.207 |
